# Ferroptosis response segregates small cell lung cancer (SCLC) neuroendocrine subtypes

**DOI:** 10.1101/2020.07.11.198408

**Authors:** Christina M. Bebber, Emily S. Thomas, Zhiyi Chen, Jenny Stroh, Ariadne Androulidaki, Anna Schmitt, Michaela N. Höhne, Lukas Stüker, Cleidson de Pádua Alves, Armin Khonsari, Marcel A. Dammert, Fatma Parmaksiz, Filippo Beleggia, Martin L. Sos, Jan Riemer, Julie George, Susanne Brodesser, Roman K. Thomas, H. Christian Reinhardt, Silvia von Karstedt

## Abstract

Bi-allelic loss of *TP53* and *RB1* in treatment-naïve small cell lung cancer (SCLC) suggests strong selective pressure to inactivate regulated cell death pathways prior to therapy. Yet, which regulated cell death pathways remain available in treatment-naïve SCLC is unknown. Here, through systemic analysis of cell death pathway availability, we identify non-neuroendocrine (NE) and NE SCLC subtypes to segregate by their response to ferroptosis, a recently described iron-dependent type of regulated necrosis. While we identify that in treatment-naïve SCLC extrinsic apoptosis and necroptosis are incapacitated, we find non-NE SCLC to be exquisitely sensitive to ferroptosis induced through pharmacological and genetic means. Mechanistically, non-NE SCLC as opposed to NE SCLC presents with an oxygenated lipidome priming non-NE SCLC for ferroptosis. ASCL1+ NE SCLC, in turn, is resistant to ferroptosis but acquires selective addiction to the thioredoxin (TRX) anti-oxidant pathway. Importantly, co-cultures mimicking non-NE/NE intratumoral heterogeneity selectively deplete non-NE populations upon induction of ferroptosis while eliminating NE cell populations only upon TRX pathway. As a consequence, combined induction of ferroptosis and inhibition of the TRX pathway broadly kills established non-NE and NE tumors in xenografts and genetically engineered mouse models of SCLC. Moreover, patient-derived treatment-naïve and refractory NE SCLC models are selectively killed via this regime. In SCLC, combined low expression of GPX4 and TRX reductase 1 (TXNRD1) identifies a patient subset with drastically improved overall survival. These data identify ferroptosis as an SCLC subtype-specific vulnerability and suggest repurposing ferroptosis induction with TRX pathway inhibition to specifically address intratumoral NE/non-NE heterogeneity in SCLC.

**One Sentence Summary:** The SCLC non-neuroendocrine subtype is sensitive to ferroptosis

## INTRODUCTION

Treatment of small cell lung cancer (SCLC) has remained a major clinical challenge, due to rapid relapse *(1)* as well as tumor heterogeneity caused by tumorigenesis from distinct cells-of-origin *(2)*. Moreover, intratumoral heterogeneity is exacerbated by standard-of-care treatment (Stewart et al. 2020, *nature cancer*) making SCLC an ultimately deadly disease. Unlike lung adenocarcinoma, SCLC rarely presents with targetable oncogenic driver mutations and is instead characterized by loss of functional p53, Rb1 and a high tumor mutational burden (TMB) *(3, 4)*. Due to the fact that SCLC presents with high TMB, a marker of potential immunotherapy response *(5)*, clinical trials involving immune checkpoint blockade have been initiated for SCLC. Although these trials have shown encouraging results *(6)*, they underperformed given the extent of response expected based on the very high TMB seen in SCLC *(7)*. However, a much-neglected fact is that high TMB may increase constitutive immune editing prior to treatment and thereby may select against pathways required for subsequent immunotherapy to function. Interestingly, caspase 8 expression, a pre-requisite for tumor cells to undergo extrinsic apoptosis triggered by immune effector cells *(8)*, was shown to be transcriptionally silenced in SCLC cell lines *(9, 10)* suggesting functional inactivation of the extrinsic apoptosis pathway in SCLC. Caspase 8 deficiency in mice causes embryonic lethality due to aberrant activation of necroptosis, a non-apoptotic form of cell death driven by receptor-interacting kinase 1 (RIPK1) and RIPK3 *(11, 12)*. Thereby, inactivation of extrinsic apoptosis can in principle cross-activate necroptosis. Interestingly, induction of ferroptosis-a recently coined type of regulated necrosis-was shown to selectively kill KRAS-driven pancreatic and lung cancer representing a novel therapeutic vulnerability in these cancers *(13, 14)*. Yet, whether SCLC, which is almost never KRAS-mutated, might respond to induction of ferroptosis has remained unexplored. Ferroptosis results from an irreparable lipid peroxidation chain reaction within cellular membranes fueled by radical formation *(15)*. Initial oxidation of membrane lipids is facilitated by divalent iron to promote a Fenton reaction generating hydroxylradicals which react with polyunsaturated fatty acids (PUFAs) *(16)*. During ferroptosis, PUFAs belonging to the arachidonic acid (AA)- and adrenic acid (AdA)-containing phosphatidylethanolamine (PE) lipid species are specifically oxidized *(17)*. Acyl-CoA synthetase long-chain family member 4 (ACSL4) generates this target lipid pool and its expression is therefore an essential prerequisite for cells to undergo ferroptosis *(18)*. *Vice versa*, Glutathione peroxidase 4 (GPX4) constitutively reduces accumulating lipid hydroperoxides thereby protecting cells from ferroptosis *(19, 20)*. GPX4 requires glutathione (GSH) as an electron donor. One important route of GSH synthesis in many cells is coupled to the availability of intracellular cysteine which can be generated from cystine imported from the extracellular space via the cystine/glutamate antiporter System xc-. Consequently, inhibition of its small subunit xCT (SLC7A11) by the small molecule erastin results in cystine depletion and ferroptosis *(21)*. Moreover, independently of GSH, ferroptosis suppressor protein 1 (FSP1, formerly AIFM2) catalyzes the production of the lipophilic radical-trapping agent ubiquinol thereby protecting several cancer entities including lung cancer from ferroptosis *(22, 23)*. Interestingly, cancer cells with acquired resistance to the HER1/HER2 tyrosinkinase inhibitor Lapatinib, so called persister cells, as well as cancer cells with a mesenchymal expression profile develop dependency on GPX4 and thereby become vulnerable to induction of ferroptosis *(24, 25)*. Importantly, ferroptosis is independent of the molecular machinery driving apoptosis and necroptosis making ferroptosis a particularly attractive type of cell death for therapeutic applications as little to no cross resistance to other types of cell death is expected to develop (recently reviewed in *(26, 27)*)

The fact that SCLC almost inevitable relapses after standard-of-care treatment fueled by increased intratumoral heterogeneity (Stewart et al. 2020, *nature cancer*) and that first-line immune checkpoint blockade only improves median survival of SCLC by two months despite high TMB *(28)* emphasizes the urgent need to initially understand the interplay of regulated cell death pathways in this type of cancer prior to treatment to devise informed treatment strategies which take cell death pathway availability into consideration.

Here, through systematic characterization of regulated cell death pathway availability in treatment-naïve SCLC, we find broad inactivation of the extrinsic apoptosis and necroptosis pathway while we identify non-neuroendocrine (NE) and NE SCLC subtypes to mechanistically segregate by ferroptosis response. NE SCLC instead acquires selective addition to the thioredoxin (TRX) pathway. Our data identify ferroptosis as a non-NE subtype-specific vulnerability in SCLC and suggest repurposing of combined induction of ferroptosis and TRX pathway inhibition as a strategy to address non-NE/NE SCLC intratumoral heterogeneity.

## RESULTS

### Treatment-naïve SCLC presents with inactivation of regulated cell death pathways

To profile cell death pathway availability in treatment-naïve SCLC, we analyzed RNA expression levels of a mostly treatment-naïve SCLC patient cohort in comparison with normal lung tissue samples of SCLC patients. SCLC primary patient tissue showed a strong downregulation of genes involved in the extrinsic apoptosis pathway (Figure 1A), such as Tumor necrosis factor-related apoptosis-inducing ligand (TRAIL, gene name TNFSF10), caspase 8 (CASP8) and CD95L (FASLG) *(8)* (Figure 1D) confirming prior observations in cell lines *(9)*. Whereas the essential extrinsic apoptosis adaptor Fas-associated protein with death domain (FADD) was slightly upregulated in SCLC, potentially facilitating non-apoptotic gene induction *(29, 30)*, the downstream essential effector caspase 8 was strongly downregulated in approximately 80% of SCLC specimens (Figure 1E), suggesting the extrinsic apoptosis pathway to be incapacitated prior to therapy. Indeed, a representative panel of human SCLC cell lines (n=7) derived prior and post-therapy *(31, 32)* was almost uniformly resistant to extrinsic apoptosis triggered by TRAIL which killed a TRAIL-sensitive control non-small cell lung cancer (NSCLC) cell (Figure 1F). Downregulation of caspase 8, can enable unrestricted activation of the necroptosis pathway through relieving caspase 8-imposed negative regulation RIPK1 and 3 *(11, 33)* (Figure 1B). Therefore, we next performed a cluster analysis of necroptosis pathway components (Figure 1G). Yet, while RIPK3 was slightly upregulated hinting at potential tumor-promoting functions *(34, 35)*, expression of the downstream essential necroptosis effector mixed lineage kinase domain like pseudokinase (MLKL) *(36–38)* was strongly downregulated in SCLC, as compared to normal lung (Figure 1H). Accordingly, SCLC cell lines were resistant to experimental induction of necroptosis by Tumor necrosis factor (TNF), zVAD and smac mimetic (TZS), in contrast to mouse embryonic fibroblast (MEFs) control cells which readily underwent necroptosis upon TZS stimulation, which was blocked by co-treatment with the RIPK1 inhibitor Necrostatin1s (Nec-1) (Figure 1I). These data suggest that strong selective pressure may already be present in treatment-naïve SCLC to incapacitate both, the extrinsic apoptosis and necroptosis pathways.

**Figure 1.**
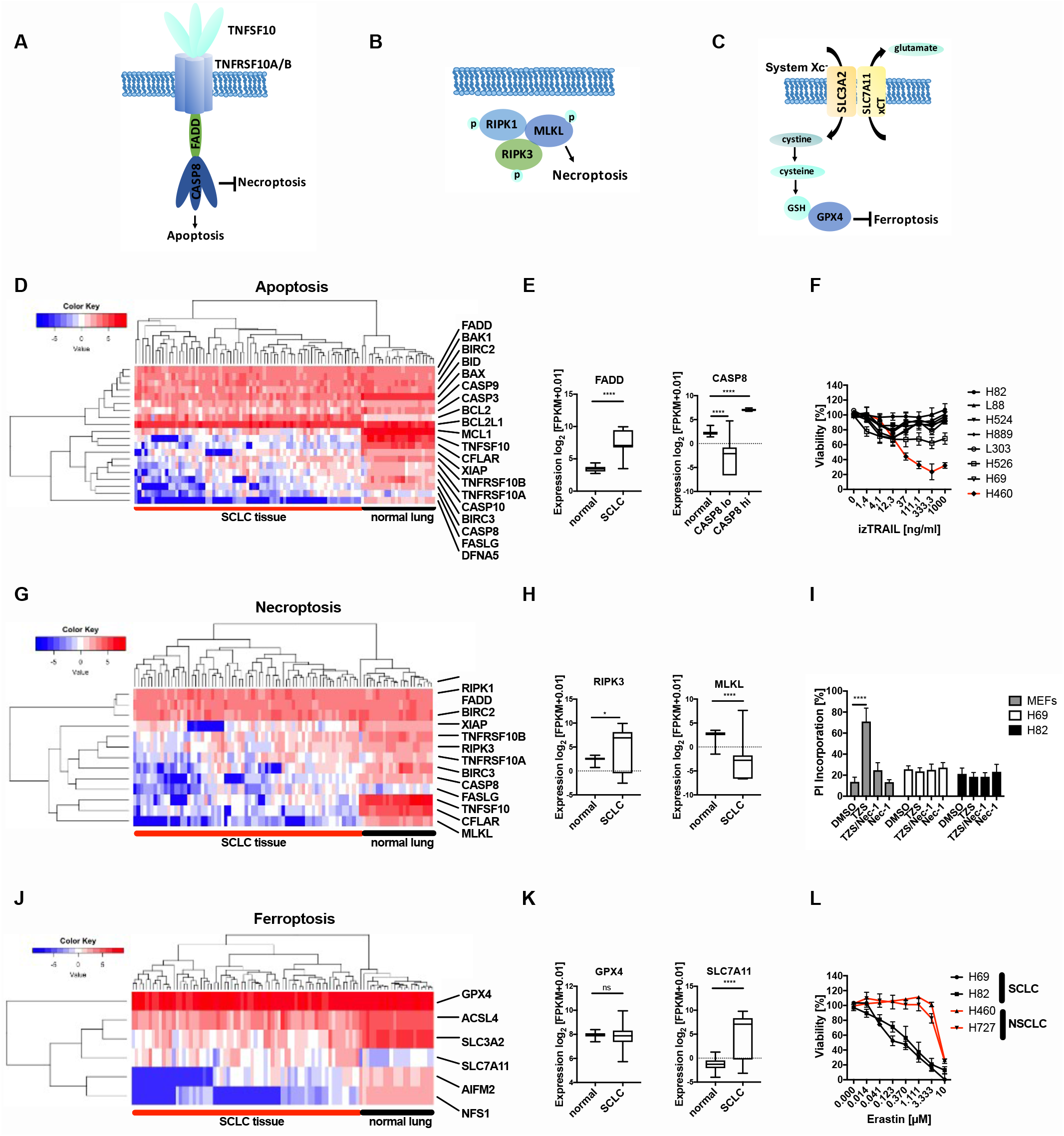
Regulated cell death pathways are counter-selected in treatment-naïve SCLC. (**A**) schematic view of genes involved in extrinsic apoptosis, (**B**) necroptosis and (**C**) ferroptosis. (**D**, **E**, **G**, **H**, **J**, **K**) RNA-seq expression data in FPKM (fragments per kilobase of exon model per million reads mapped) from normal lung *(70)* (n=22) and mostly chemo-naïve SCLC patient samples *(3)* (n=67) were log2 transformed (+0.01) and plotted for relative expression of genes involved in extrinsic apoptosis (**D**, **E**) necroptosis (**G**, **H)** and ferroptosis (**J**, **K)** boxplot center line, mean; box limits, upper and lower quartile; whiskers min. to max. (**F**) The indicated human SCLC cell lines (n=7) and NSCLC line (H460) were treated with human izTRAIL for 24 h, cell viability was determined by Cell Titer Blue. (**I**) Cells were treated with TNF (T) [10 ng/ml]/zVAD (Z) [20 μM]/Smac mimetic (S) Birinapant [1 μM] for 24 h, cell death was quantified by propidium iodide (PI) uptake and flow cytometry. (**L**) Cells were treated with erastin at indicated concentrations for 24 h, cell viability was determined by Cell Titer Blue. Data are means +/− SEM of three independent experiments wherever not indicated otherwise.

Recently, ferroptosis, an iron-dependent form of regulated necrosis was described *(39, 40)*. Importantly, ferroptosis is independent of the molecular machinery driving apoptosis or necroptosis (Figure 1C). Therefore, we next assessed expression of ferroptosis regulators in SCLC patient samples. Strikingly, xCT was strongly upregulated in SCLC and only expressed at very low levels in normal lung (Figure 1J, K). GPX4 was highly and comparably expressed in SCLC and normal lung. Moreover, in human SCLC cell lines, protein expression of both xCT and GPX4 could be validated (Figure S1A). Elevated xCT expression might therefore represent a selective advantage for SCLC - as opposed to normal lung - exposing a potential SCLC-selective therapeutic opportunity. Confirming this notion, treatment of SCLC cells, but much less of NSCLC cells, with the xCT inhibitor erastin resulted in dose-dependent cytotoxicity (Figure 1L). Moreover, when directly comparing erastin sensitivity of murine SCLC cells from genetically-engineered mouse models for SCLC *(41)* and NSCLC *(42)*, again SCLC cells were overall more sensitive to erastin-induced cytotoxicity (Figure S1B). In addition, viability data taken from an erastin screen in 117 cancer cell lines *(20)* independently confirmed increased erastin sensitivity of SCLC, over comparable NSCLC cell lines (Figure S1C). Taken together, we find that SCLC has evolved to present with cell death resistance on multiple levels prior to treatment, i.e. against extrinsic apoptosis, necroptosis and ferroptosis. However, since ferroptosis escape in SCLC, unlike escape from other cell death pathways, involves the upregulation of protective and targetable proteins, rather than the loss of agonists, we next aimed to mechanistically validate the induction of ferroptosis in SCLC.

### SCLC is vulnerable to the induction of ferroptotic cell death

To validate whether SCLC was vulnerable to ferroptotic cell death in general, murine SCLC cells were treated with the GPX4 small molecule inhibitor RSL3, known to trigger ferroptosis *(20)* with or without the ferroptosis-selective lipophilic radical scavenger Ferrostatin-1 (Fer-1) *(39)*. Indeed, RSL3 effectively abrogated clonogenic survival of adherent murine SCLC lines, which could be restored with Fer-1 co-treatment (Fig. 2A). Moreover, cell death induced by erastin or two structurally distinct small molecule inhibitors against GPX4, RSL3 *(43)* and ML210 *(44)*, could be rescued by co-incubation with Fer-1 (Fig. 2B, C S2A). Indicating requirement for free iron in cell death execution, co-incubation with the iron-scavenger deferoxamine (DFO), equally rescued cell death induced by erastin or GPX4 inhibition (Fig. 2D, E). Lastly, cell death induced by erastin or RSL3 in murine SCLC cell lines could also be rescued by Fer-1 (Fig. S2B, C). While GPX4 is central to protecting a variety of cells from ferroptosis, in lung adenocarcinoma cells, GPX4 deletion is not sufficient to induce ferroptosis *(23)*. Therefore, to determine whether or not GPX4 is sufficient to protect SCLC from ferroptosis, we generated CRISPR/Cas9-mediated control or GPX4 knockout (KO) SCLC lines. Supporting a vital function for GPX4 in preventing toxic build-up of lipid reactive oxygen species (ROS) in SCLC, these cells could only be generated and cultured in the presence of Fer-1 in the media. Consequently, upon Fer-1 withdrawal, cells with GPX4-targeting gRNAs selectively died (Fig. 2F, Fig. S2D). GPX4 KO was confirmed on protein level in the presence of Fer-1 (Fig. 2G). In addition, GPX4 KO SCLC cells presented with lipid ROS accumulation upon Fer-1 withdrawal, indicative of the induction of ferroptotic cell death *(39)* (Fig. 2H, I, Fig. S2E). Collectively, these data support a requirement for free iron and lipid radicals in cell death execution upon pharmacological induction of ferroptosis and appoint GPX4 as the central player sufficient to prevent lipid peroxidation and ferroptosis in SCLC.

**Figure 2.**
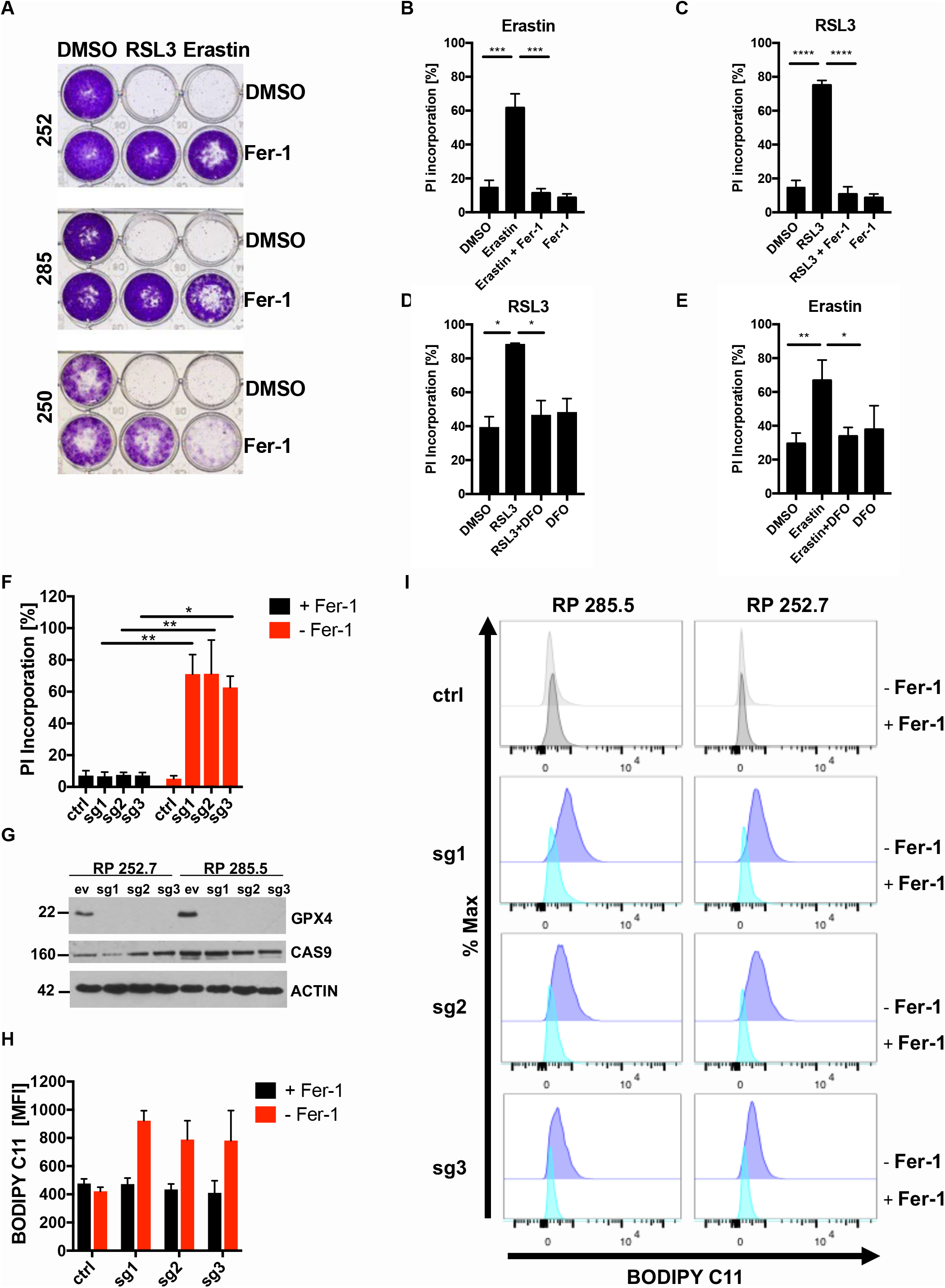
GPX4 expression is sufficient to protect SCLC from ferroptosis. (**A**) the indicated murine SCLC lines (n=3) were treated either with DMSO, RSL3 [1 μM], erastin [10 μM] alone or in combination with Ferrostatin-1 (Fer-1) [5 μM] for 24 h. Cells were washed, cultured for 5 days for recovery and stained with crystal violet. (**B, C, D, E**) Human H82 cells were treated as indicated: erastin [10 μM] +/− Ferrostatin-1 (Fer-1, 5 μM) +/− Deferoxamine (DFO) [100 μM], RSL3 [1 μM] +/− Fer-1 +/− DFO for 24 h. Cell death was determined by propidium iodide (PI) uptake and flow cytometry. (**F**) RP285.5 murine SCLC cells stably expressing Cas9 and the indicated control or gRNAs targeting GPX4 were cultured with or without Ferrostatin-1 (Fer-1) for 24 h. Cell death was quantified by propidium iodide (PI) uptake and flow cytometry. (**G**) protein extracts were obtained from cells as in (**F**) cultured in the presence of Fer-1 [5 μM]. (**H, I**) RP285.5 murine SCLC cells as in (**F**) were cultured in the presence or withdrawal of Fer-1 [5 μM] for 5 h and stained for lipid ROS accumulation using BODIPY C11. Cells were analyzed by flow cytometry and mean fluorescent intensity (MFI) was quantified. Data are means +/− SEM of three independent experiments in each individual cell line or representative images or histograms were applicable.

### Non-NE SCLC is exquisitely sensitive to lipid peroxidation

SCLC can be subdivided into several molecular subtypes based on patterns of expression of neuroendocrine (NE) differentiation markers, which differ in their cell biology *(45)*, cell of origin or arise as a consequence of intratumoral plasticity *(46)*. Therefore, we next aimed to understand whether ferroptosis sensitivity was a common feature of all molecular subtypes of SCLC. To address this, we tested ferroptosis sensitivity in a larger panel (n=8) of human SCLC lines representative of described SCLC molecular subtypes *(1)*. Interestingly, we noted that this panel divided into subsets of responders (black) and non-responders (grey) in response to all inducers of ferroptosis tested (Figure 3A, B and S3A). This separation was also reflected in failure to accumulate lipid ROS upon GPX4 inhibition in non-responders (Figure 3C). A division into ferroptosis responders and weak-responders with impaired accumulation of lipid ROS upon ferroptosis induction was equally observed in a panel of murine SCLC lines, albeit less pronounced (Figure S3B-D). Of note, this divided response pattern was unique to ferroptosis, as the same murine and human SCLC cell line panels did not show a bipartite response to cisplatin or etoposide (Figure S4A-D). Cisplatin, as well as etoposide, instead led to caspase-dependent cell death as pan-caspase inhibitor zVAD partially reverted cell death induction (Figure S4E, F).

**Figure 3.**
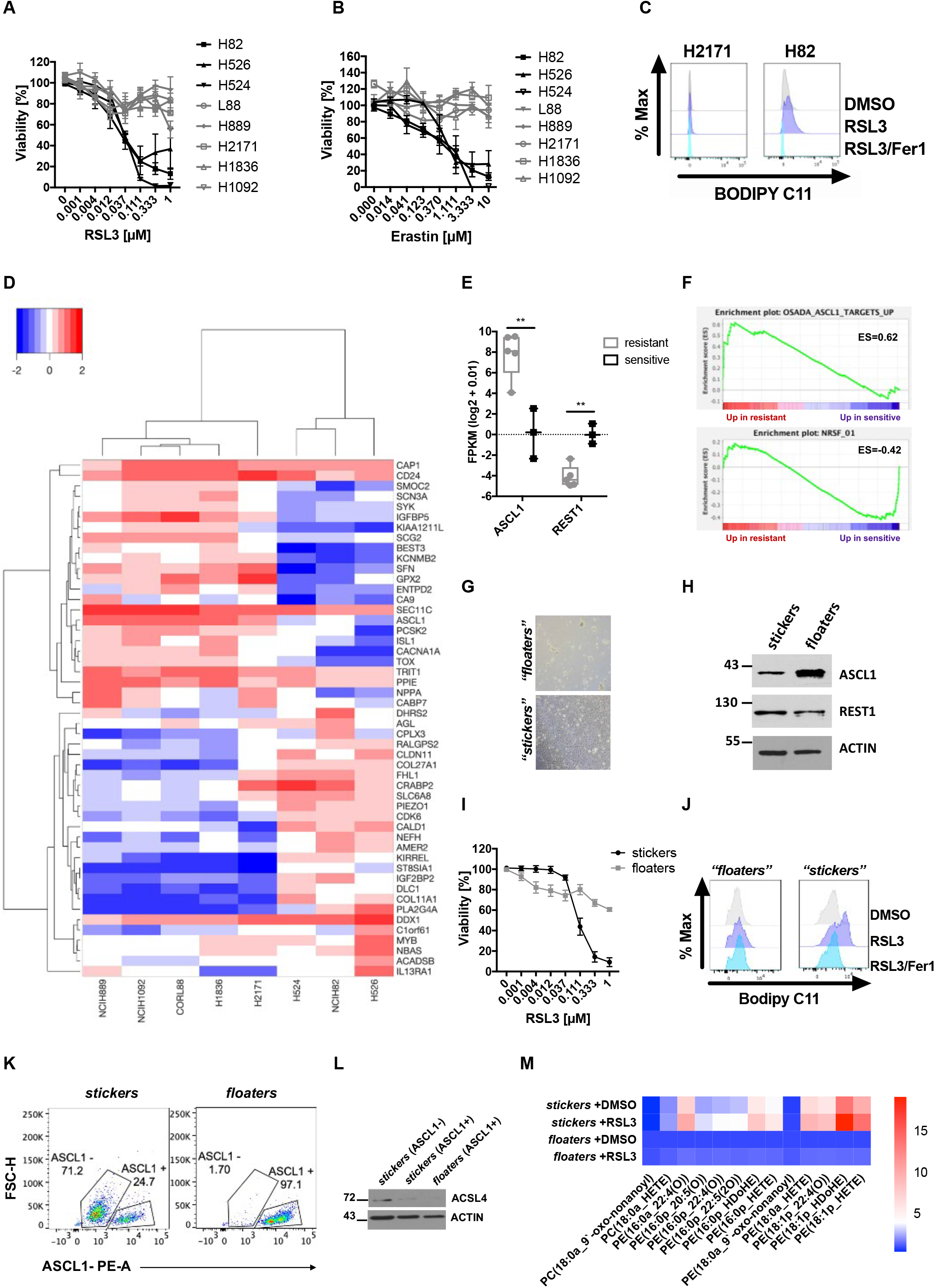
SCLC neuroendocrine subtypes segregate by ferroptosis response. (**A**, **B**) Human SCLC cell lines (n=8) were treated as indicated for 24 h, cell viability was determined by Cell Titer Blue. (**C**) the indicated human SCLC cells were treated with DMSO, RSL3 [100 nM] or RSL3/Fer-1 [5 μM] for 5 h and stained for lipid ROS accumulation using BODIPY C11. Cells were analyzed by flow cytometry. (**D**) RNA-seq data *(47)* of human SCLC lines (sensitive n=3, H524, NCIH82, H526; resistant n=5, NCIH889, NCIH1092, CORL88, H1836, H2171) were analyzed for differential expression between responders and non-responders, heatmap represents hierarchical clustering of FPKM (log2+0.01) of the 50 most differentially expressed genes. (**E**) ASCL1 and REST1 expression (FPKM (log2+0.01) comparing sensitive and resistant cells is plotted, boxplot center line, mean; box limits, upper and lower quartile; whiskers min. to max. (**F**) Gene set enrichment (GSEA) of a ranked list from ferroptosis sensitive and resistant cells was performed. (**G**) *stickers* and *floaters* were cultured separately; representative images were taken at 200x magnification. (**H**) western blot of SCLC subtype marker expression in manually separated *stickers* and *floaters*. (**I**) *stickers* and *floaters* were cultured separately and treated with RSL3 at the indicated concentrations for 24 h, cell viability was determined by Cell Titer Blue. (**J**) *stickers* and *floaters* were cultured separately and treated with DMSO, RSL3 [1 μM] +/− Fer-1 [5 μM] for 5 h and stained for lipid ROS accumulation using BODIPY C11 and analyzed by flow cytometry; mean fluorescent intensity (MFI) was quantified. (**K**) gating strategy for ASCL1 high *stickers* and *floaters* FACS sorting. (**L**) *stickers* and *floaters* were FACS-sorted by ASCL1 expression and ACSL4 protein levels were detected by Western blot. (**M**) Heatmap showing the representation of mono-oxidized phospholipid species (PE, phosphatidylethanolamine; PC, phosphatidylcholine) in *stickers* as compared to *floaters* treated with either DMSO or RSL3 [1 μM] for 5 h and then subjected to lipidomics. Samples for each condition (n=5) were averaged and normalized to the cell number (2.5×10^6^). Each lipid species was normalized to levels detected in *floaters* +DMSO. One representative out of two independent experiments is shown. Data are means +/− SEM of three independent experiments or representative images if not indicated otherwise.

To identify markers and mechanisms of ferroptosis sensitivity and/or resistance in SCLC responders and non-responders, we analyzed mRNA expression patterns of human SCLC cells *(47)*, comparing ferroptosis responders and non-responders which clustered into two distinct groups (Fig. 3D). Interestingly, while non-responders showed increased expression of the transcription factor Achaete-Scute Homolog 1 (ASCL1), responders instead expressed elevated levels of RE1-silencing transcription factor (REST1, also known as neuron-restrictive silencing factor NRSF) (Fig. 3E). In line with this, gene set enrichment analysis (GSEA) revealed that upregulated ASCL1 target genes *(48)* were enriched in non-responders while a REST1 signature was enriched in ferroptosis sensitive cells (Fig. 3F). While ASCL1 is known to promote NE differentiation in SCLC, REST1 suppresses this process. Intriguingly, SCLC cells can spontaneously transdifferentiate from non-NE (REST1+, ASCL1−) to an NE-ASCL1+ (NE) state phenotypically marked by adherent growth (“*stickers*”) and growth in suspension (“*floaters*”), respectively *(45)*. Making use of this system of isogenic spontaneous NE differentiation, we obtained SCLC cells which spontaneously segregated into *stickers* and *floaters* (Fig. 3G) and first validated expression of ASCL1 as NE marker and REST1 as a non-NE marker in this cellular system (Fig. 3H). Strikingly, isogenic NE *floaters* were indeed more resistant to ferroptosis triggered by GPX4 inhibition, or erastin, while non-NE *stickers* were sensitive (Fig. 3I and S5A). Consequently, *floaters* unlike *stickers* showed impaired accumulation of lipid ROS upon GPX4 inhibition (Fig. 3J, S5B). Of note, this segregation of cell death sensitivity was specific to ferroptosis, as *stickers* and *floaters* responded equally to cisplatin or etoposide treatment (Figure S5C, D). Moreover, cell adherent or non-adherent states in general did not determine ferroptosis sensitivity or resistance, as all human cell lines tested, including responders and non-responders, grow in suspension (Fig. 3A). To validate ferroptosis response segregation by NE and non-NE differentiation in an independent experimental set-up, we made use of SCLC cells with cMyc expression induced from the endogenous locus via CRISPR activation (CRISPRa) *(49)* as cMyc was shown to promote transition to a non-NE SCLC phenotype *(46, 47)*. Indeed, cMyc expression sensitized SCLC cells to ferroptosis induced by GPX4 inhibition (Figure S5E, F).

To determine factors mediating ferroptosis sensitivity in non-NE SCLC, we sorted *stickers* and *floaters* by ASCL1 expression using flow cytometry. We noted that within *stickers* there was an emerging ASCL1^high^ population whereas all *floaters* were ASCL1^high^ (Fig. 3K). Strikingly, protein levels of ACSL4, which is essential for esterification of arachidonic- and adrenic acid to generate PE target lipid species specifically peroxidized during ferroptosis *(17)*, inversely correlated with ASCL1 expression in *stickers* and *floaters* (Fig. 3L). Moreover, in the NIH SCLC cell line panel, ASCL1 mRNA expression inversely correlated with ACSL4 expression (Fig. S5G) proposing increased availability of PE peroxidation target lipids to facilitate ferroptosis in non-NE SCLC. Therefore, we next determined how NE differentiation in isogenic *stickers* as compared to *floaters* may affect the oxidized lipidome upon induction of ferroptosis using mass spectrometry. While basal ferroptosis-specific phospholipid peroxidation products were indeed elevated in *stickers* and further increased upon RSL3 treatment, both, the basal amount and the specific induction of these products was markedly decreased in *floaters* (Fig. 3M and Supplementary Table 1). Therefore, we find non-NE SCLC to be characterized by high ACSL4 expression, a ferroptosis-prone lipidome and exquisite sensitivity to ferroptosis.

### NE SCLC is defined by selective addiction to the TRX anti-oxidant pathway

While we found non-NE SCLC to be ferroptosis sensitive, NE SCLC was highly resistant. In order to determine whether NE SCLC may have acquired ferroptosis resistance through upregulating GSH and thereby GPX4 activity, we measured cellular concentrations of GSH and its oxidized form GSSG. Surprisingly and contrary to expectations, NE *floaters* and human and murine resistant NE cells in fact presented with lower basal levels of GSH whilst levels of reduced GSSG were comparable with those in responders and *stickers* (Fig. S6A, B). These data indicated that GSH synthesis but not recovery is specifically repressed in ASCL1-expressing NE SCLC. In support of this, inducible expression of ASCL1 suppressed expression of glutamate-cysteine ligase catalytic subunit (GCLC) an essential enzyme in the synthesis of GSH (Fig. S6C). Consequently, inducible expression of ASCL1 also suppressed cellular amounts of GSH (Fig. S6D). Therefore, ASCL1 expression selectively lowers the cellular redox potential endowed by cellular GSH, strongly suggesting that NE SCLC must adopt alternative means of anti-oxidant defense whose targeting may represent an alternative vulnerability in NE SCLC.

Loss of the GSH antioxidant pathway is known to be compensated for by the thioredoxin (TRX) antioxidant pathway *(50, 51)*. As part of the thioredoxin (TRX) antioxidant pathway, thioredoxin reductase 1 (TrR1, gene name TXNRD1) reduces oxidized TRX which can then function to reduce other oxidized cellular substrates. Reduced TRX can in turn be sequestered by TXNIP limiting TRX availability for antioxidant defense (Fig. 4A). To understand whether NE SCLC may regulate TRX pathway components to compensate for low cellular GSH, we monitored their protein expression comparing human non-NE with NE cells as well as *stickers* with *floaters*. Interestingly, while TrR1 and TrR2 were upregulated only in human NE SCLC cells, expression of TXNIP was consistently increased in human and murine NE cells suggesting NE SCLC cells to also experience a limited availability TRX-mediated anti-oxidant defense (Fig. 4B and S6E). Indeed, treatment with the TrR1 and TrR2 inhibitor Auranofin - clinically approved for the treatment of rheumatoid arthritis-led to a quicker time-dependent loss of fully-reduced TRX in NE as compared to non-NE SCLC cells supporting pre-existing inhibition of the TRX pathway in NE-A SCLC (Fig. 4C and S6F). Strikingly, this translated into selective sensitivity to Auranofin-induced cell death in NE SCLC while non-NE SCLC was more resistant to Auranofin (Fig. 4D). The same NE-selective effect could be observed for two other structurally distinct inhibitors of TrR1 (PX 12 and D9) (Figure S6G, H). *Vice versa*, the same set of cell lines showed an inverse response pattern to ferroptosis induced by the clinically advanced GCLC inhibitor buthionine sulfoximine (BSO) (Fig. 4E). While BSO triggered ferrostatin-1 blockable ferroptosis in non-NE *stickers*, Auranofin-induced cell death in NE *floaters* was non-ferroptotic (Fig. 4F). Similarly, cystine starvation induced ferroptosis in *stickers*, but not in *floaters* (Figure S6I). Therefore, NE SCLC is selectively addicted to the TRX anti-oxidant pathway while non-NE is sensitive to ferroptosis, a concept which holds true in isogenic cells spontaneously undergoing non-NE/NE plasticity.

**Figure 4.**
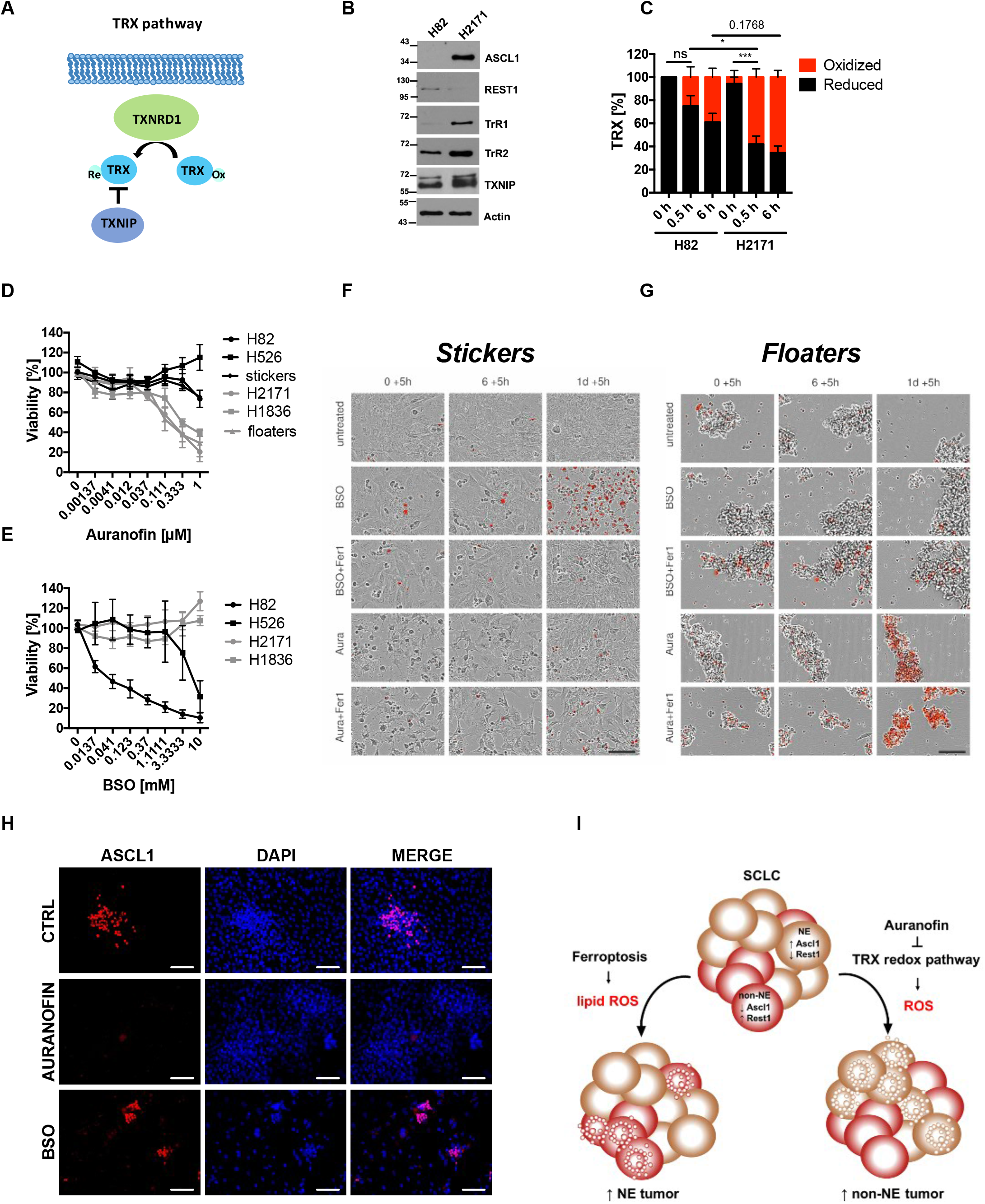
Neuroendocrine SCLC presents with TRX pathway addiction. (**A**) schematic view of genes involved in the TRX antioxidant pathway. (**B**) H82 and H2171 cells were lysed and TRX pathway component expression was detected by Western blot. (**C**) indicated cells were treated with Auranofin [1 μM] for the indicated times and subjected to redox shift assays. Densitometrical quantification of TRX redox forms is shown. (**D**) indicated cells were treated with Auranofin for 24 h, cell viability was determined by Cell Titer Blue. (**E**) indicated cells were treated with BSO for 24 h, cell viability was determined by Cell Titer Blue. (**F**, **G**) RP181.5 manually separated *stickers* and *floaters* were treated with Ferrostatin-1 [5 μM] for 2 h prior to adding DMSO, BSO [10 mM] or Auranofin [1 μM] for an additional 24h. DRAQ7 [0,1 μM] (red color in image) was added to all wells to visualize dead cells. Images were acquired every 5 h using the IncuCyte S3 bioimaging platform. Scale bar = 100 μm. Data are means +/− SEM of three independent experiments or representative images where applicable. (**H**) RP181.5 *stickers* were treated with either DMSO, Auranofin [500 nM] or BSO [500 μM] for 96 h and then fixed and stained for ASCL1 (red) and counterstained with DAPI (blue). Scale bar = 100 μm. (**I**) schematic view of SCLC NE subtype ferroptosis vulnerability and its effect in heterogeneous tumors. Data are means +/− SEM of three independent experiments or representative images where applicable.

SCLC tumors are known to present with intratumoral heterogeneity in respect to individual cellular NE differentiation *(45, 47)*. Moreover, this NE subtype intratumoral heterogeneity can originate from plasticity *(46)*. Therefore, to mimic a situation of NE intratumoral heterogeneity, we next determined how induction of either ferroptosis or TRX pathway inhibition would affect an isogenic mixed ASCL1^+^/ASCL^−^ (non-NE/NE) culture. For this, we made use of the fact that the manually separated *stickers* culture already contains a subpopulation of cells in the process of NE transdifferentiation with high expression of ASCL1 (see also Fig. 3K). Strikingly, while a morphologically distinct subpopulation of SCLC cells with bright nuclear ASCL1 staining could be detected by microscopy in control treated cells, this population was selectively depleted upon treatment with Auranofin which did not affect numbers of ASCL1^−^ cells as visualized by DAPI. *Vice versa*, induction of ferroptosis using BSO led to a strong relative enrichment in ASCL1^+^ cells while the majority of ASCL1^−^ cells was killed (Fig. 4H). Together, these data suggest a model for homogeneous tumors in which ferroptosis induction kills non-NE SCLC while TRX pathway inhibition kills NE SCLC. In heterogeneous SCLC tumors, however, our data suggest that fractional killing via ferroptosis will select for NE SCLC over time while, *vice versa*, TRX pathway inhibition might enrich tumors for non-NE SCLC cells (Fig. 4I). Therefore, we next evaluated efficacy of combining ferroptosis induction *in vivo* with TRX pathway inhibition in SCLC non-NE and NE subtypes.

### Combining ferroptosis induction with TRX pathway inhibition demonstrates broad therapeutic efficacy across SCLC NE subtypes *in vivo* and serves as prognostic marker set in human SCLC

To circumvent SCLC subtype tumor plasticity under either ferroptosis induction or TRX redox pathway inhibition, we aimed to simultaneously induce ferroptosis together with inhibition of the TRX pathway *in vivo*. Although several efforts are underway to develop selective GPX4 inhibitors with improved bioavailability for clinical applications, these do not meet required pharmacokinetics yet *(26, 27)*. Therefore, we made use of the fact that the clinically advanced GCLC inhibitor BSO induced lipid ROS-dependent ferroptosis in non-NE SCLC cells (Fig. 4F). Moreover, to equally repurpose a clinically applied inhibitor for the TRX pathway to facilitate rapid clinical translation for the treatment of SCLC, we used Auranofin *in vivo*. To now test non-NE and NE SCLC *in vivo* sensitivity, representative NE and non-NE SCLC cells were subcutaneously implanted into nude mice. Upon detection of palpable tumors, mice were randomized by comparable tumor volume (Figure S7A, B) and treated with either vehicle or well tolerated combined BSO/Auranofin (combo) (Figure S7C, D). Strikingly, we observed a very clear and significant response in established NE and non-NE SCLC tumors *in vivo* (Fig. 5A, B). These data suggested that SCLC may be a particularly sensitive entity for this strategy due to its NE subtypes being addicted to mutually exclusive anti-oxidant pathways. Cell death induced by this treatment combination in SCLC was partially dependent of lipid ROS and, thereby, ferroptotic and partially dependent on general ROS (Figure S7E, F). Confirming target specificity of this regime in SCLC, only combined siRNA-mediated suppression of GCLC, TrR1 and TrR2, led to an equal extent of specific cell death induction in *stickers* and *floaters* (Figure S7G). Importantly, the lipid ROS generation byproduct malondialdehyde (MDA), recently validated as a specific *in vivo* marker for ferroptosis induction*(53)*, was increased in combo-treated tumors of both subtypes (Fig. 5C).

**Figure 5.**
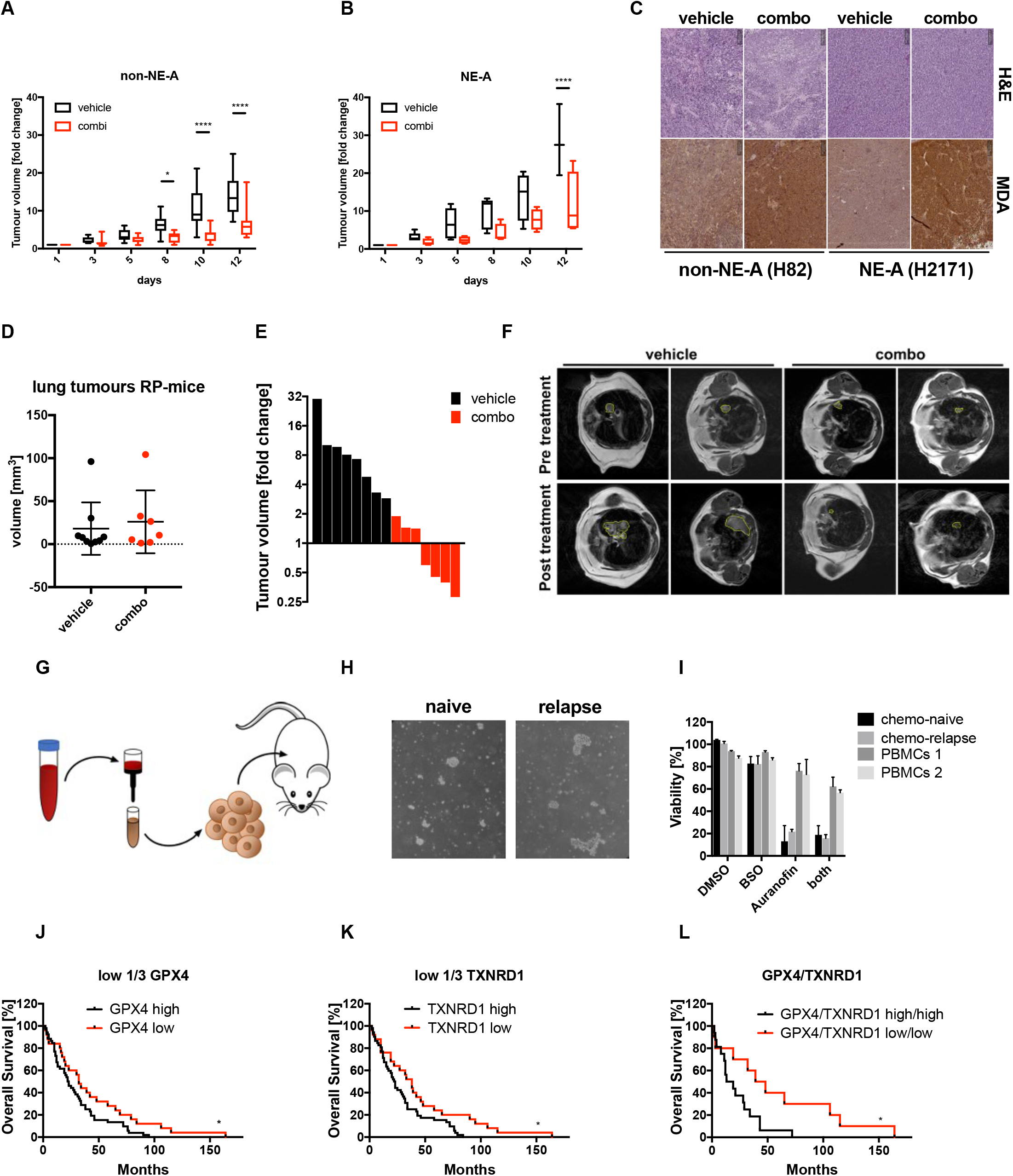
Combined ferroptosis induction and TRX pathway inhibition demonstrates broad anti-tumor activity across SCLC NE subtypes *in vivo* and serves as prognostic marker set in human SCLC. (**A**, **B**) 8-weeks old male nude mice were injected with 1.5×10^6^ H82 (**A**) and H2171 (**B**) cells into flanks. Once palpable, tumors were treated either with vehicle (H82, n=9; H2171 n=4) or combined BSO [5 mM] in the drinking water and Auranofin 3x per week i.p. [2.5 mg/kg] (H82, n=11, H2171 n=4) for two consecutive weeks. Fold change of initial tumor size is shown. (**C**) Sections from paraffin-embedded tumors of vehicle or combo-treated mice were stained by H&E or for MDA. Representative images are shown, scale bar=200 μm. Data are means +/− SEM of three independent experiments or representative images where applicable. (**D**) lung tumor volumes were quantified by Horos software using MRI scans. Mice were randomized to obtain two groups with equal mean tumor volume (n=8 for vehicle, n=7 for combo). **(E)** tumor-bearing RP-mice were treated either with vehicle (n=8) or combined BSO [5 mM] in the drinking water and Auranofin 3x per week i.p. [2.5 mg/kg] (n=7) for two consecutive weeks. Fold change in tumor volumes was determined by quantifying initial tumor volume from MRI scans as compared to tumor volume at the end of the treatment cycle using Horos software. (**F**) representative MRI images pre and post treatment of mice as in (**E**). (**G**) isolation scheme of human CDXs. (**H**) Cellular morphology of human CDXs. (**I**) CDXs (n=2) or healthy donor PBMCs (n=2) were treated with DMSO, BSO [PBMCs, 500 μM; CDXs, 50 μM], Auranofin [250 nM] or BSO [PBMCs, 500 μM; CDXs, 50 μM] /Auranofin [250 nM] for 24 h, cell viability was quantified by Cell Titer Blue (CDXs) or flow cytometric quantification of propidium iodide (PI)-negative cells (PBMCs). (**J**) Kaplan-Meier survival curves for SCLC patients (n=77) *(3)* containing low (low 1/3 n=25, median survival 33 months) or high (high 2/3 n=52, median survival 22.5 months) expression of GPX4 mRNA. (**K**) as in (**J**) expression of TXNRD1 mRNA was correlated using the same cut-off (low=1/3, median survival 38 months; high 2/3, median survival 22.5 months). (**L**) Kaplan-Meier survival curves for SCLC patients with combined low or high GPX4 and TXNRD1 mRNA expression (low/low n=10, median survival 43.5 months; high/high n=16, median survival 16 months). Data are means+/− SEM were applicable (**E**).

To next validate efficacy in an immune-proficient autochthonous mouse model, we made use of an established genetically engineered mouse model (GEMM) recapitulating all relevant features of human SCLC *(41)*. In these mice, SCLC develops as a consequence of Rb1 and Tp53 co-deletion (RP-mice) upon adenoviral Cre inhalation within 9 months. Moreover, intratumoral heterogeneity is observed in these mice as tumors contain both ASCL1^high^ and ^low^ cells *(45)*. Once tumors had a detectable mean volume of 15-30 mm^3^, determined by magnetic resonance imaging (MRI), mice were randomized (Fig, 5D) and treated with combo for 2 weeks. Whereas, all vehicle control mice progressed, 4 out of 7 tumors receiving combo significantly regressed (Fig. 5E, F).

Besides the use of GEMMs, circulating tumor cells (CTCs) and CTC-derived xenotransplants (CDXs) from SCLC have proven to be a powerful tool for faithful recapitulation of patient response to chemotherapy *(54, 55)*. In order to test a potential response to combo in this human model system, we obtained CDXs from a treatment-naïve and a post-chemotherapy relapse SCLC patient (Fig. 5G, H). Both CDXs, which were confirmed to express NE markers, were sensitive to Auranofin but not ferroptosis induced by BSO whereas normal peripheral blood monocytes (PBMCs) were resistant (Fig. 5I). Importantly, when treating these cells with combo, similar levels of killing could be achieved in chemotherapy-naïve and relapse CDXs (Fig. 5I) indicating that prior chemotherapy and relapse does not impact response to combo. To lastly determine whether protection from ferroptosis via GPX4 or protection from ROS-dependent cell death via TrR1 (TXNRD1) is of prognostic value in SCLC, overall survival of patients who had undergone surgical resection *(3)* was analyzed. Interestingly, low GPX4 or low TrR1 (TXNRD1) expression, both independently correlated with improved overall survival (Fig. 5J, K). Importantly, when analyzing SCLC patients with combined low expression of GPX4 and TrR1 mimicking combo treatment, we obtained a group of patients with a drastically improved median survival time of 43.5 months as compared to 16 months median survival in the high/high group (Fig. 5L). This expression pattern did not correlate with treatments these patients received post-surgery (chemotherapy or radiation) *(3)*. Moreover, to remove survival bias originating from advanced tumor stage at diagnosis, diagnosed stage IV patients were excluded from the analysis and stage III patients were equally present in both groups (n=4 in high/high; n=3 in low/low). Importantly, survival in lung adenocarcinoma (LUAD) did not significantly segregate by GPX4, TXNRD1 or combined GPX4 and TXNRD1 expression (Figure S7E-G) suggesting a function for redox pathway plasticity in determining patient outcome to be a unique feature of SCLC. Therefore, concomitant low expression of GPX4 and TRXNRD1 serves as an independent and specific prognostic marker set for overall survival in SCLC.

In conclusion, we report that treatment-naïve SCLC exhibits signs of selection against extrinsic apoptosis and necroptosis and upregulates xCT for ferroptosis protection. While we identify the non-NE-A SCLC subtype to be exquisitely ferroptosis sensitive and present with high ACSL4 expression and a ferroptosis-prone lipidome, we find the NE-A SCLC subset to be resistant to ferroptosis but selectively vulnerable to TRX pathway inhibition. Thereby, we identify that SCLC non-NE/NE-subtypes mechanistically segregate by ferroptosis sensitivity or resistance. Importantly, heterogeneous SCLC cultures selectively deplete non-NE or NE subpopulations upon single pathway targeting. Hence, combining ferroptosis induction with TRX pathway inhibition demonstrates high therapeutic efficacy in SCLC xenografts, GEMMs, patient-derived CDXs and identifies a unique SCLC patient subset with drastically improved prognosis. Taken together, we reveal ferroptosis as an Achilles heel of non-NE SCLC and TRX pathway inhibition as a novel NE SCLC vulnerability, a concept which proposes that combined ferroptosis induction/TRX pathway inhibition may specifically tackle the problem of intratumoral NE/non-NE heterogeneity in SCLC.

## DISCUSSION

SCLC is amongst the deadliest human cancers with a characteristically high TMB (4), a feature known to positively correlate with response to immunotherapy *(7, 56)*. Although immunotherapy has resulted in unprecedented responses in advanced melanoma *(57)*, median survival of SCLC patients treated with immunotherapy was only improved by two months *(28)*. The extrinsic apoptosis pathway is preferentially triggered by activated immune effector cells for anti-tumor immune-surveillance (8). As such, intra-tumoral profiles of cytolytic immune activity involved inactivating mutations of caspase 8 along with loss of the antigen presenting machinery in various cancers *(58).* In this study, we show that the extrinsic apoptosis pathway is already disabled in treatment-naïve SCLC tissue (Fig. 1D-F). Importantly, we find that compensatory activation of the necroptosis pathway is equally eliminated at this early stage (Fig. 1F-H), inactivation of both pathways may explain the suboptimal response to immunotherapy in SCLC. Nevertheless, immunotherapy-activated CD8 T-cells were recently shown to suppress expression of xCT sensitizing tumor cells to ferroptosis *(59)*. Thereby, promotion of ferroptosis can equally be part of the cell death arsenal triggered by activated immune effector cells. Consequently, over time, tumors exposed to immune-surveillance may also upregulate proteins protecting from ferroptosis. In fact, upregulation of NFS1 which protects from ferroptosis has been observed for non-small cell lung cancer (NSCLC) in the context of an oxygenated microenvironment *(60)*. Of note, we did not observe upregulation of NFS1 in SCLC but instead xCT, suggesting cystine import for GSH synthesis to be a more important bottleneck in SCLC.

Importantly, GPX4 knockout was insufficient to induce ferroptosis in NSCLC cells which required additional deletion of FSP1 *(22, 23)*. In contrast to this, SCLC cells readily underwent ferroptosis with genetic deletion of GPX4 upon withdrawal of Fer-1, highlighting that SCLC is not additionally protected by FSP1 which may be responsible for the exquisite sensitivity observed for the non-NE subtype. Interestingly, we found NE-differentiation to determine ferroptosis resistance in SCLC (Fig. 3). NE to non-NE fate switch in SCLC is a particularly interesting feature as it is closely linked with resistance to conventional chemotherapy. Non-NE cells within tumors can share a common origin with NE cells proposing a high degree of plasticity within SCLC *(61)*. Importantly, these non-NE cells are characterized by a mesenchymal phenotype and expression profile. Of note, a mesenchymal expression profile has been identified as a ferroptosis response signature *(25)*. Moreover, remodeling of the plasma membrane during EMT leads to an increase in biosynthesis of PUFAs which are the main target of lipid peroxidation, the fatal event in the execution of ferroptosis *(62)*. The key enzyme generating the lipid target pool for lipid peroxidation is ACSL4 which we find to be expressed at higher levels in non-NE SCLC (Fig. 3L). Interestingly, Notch homolog 1 (NOTCH1) signaling has repeatedly been shown to promote expression of an EMT signature in various cancers *(63)*. In SCLC, NOTCH signaling promotes a slow growing and chemotherapy resistant non-NE cell fate via activation of REST1 *(45)*. Therefore, it is tempting to speculate whether NOTCH pathway activation and a resulting EMT may set these intrinsically chemotherapy resistant cells up to be ferroptosis responsive.

Besides its role in driving NE cell fate in SCLC, ASCL1 expression has been instrumental in neuronal reprogramming. Here, upon ASCL1 expression cells have to overcome a period of enhanced sensitivity to ferroptosis to fully transdifferentiate into neurons *(64)*. Thereby, to tolerate ASCL1 expression these cells must acquire ferroptosis resistance through selection and adaption rather than through direct ASCL1-mediated ferroptosis resistance. We propose that this may equally be the case in SCLC as acute ASCL1 overexpression did not render cells resistant to ferroptosis, yet, all ferroptosis resistant SCLC cells express ASCL1. In line with an idea of increased ferroptosis selective pressure arising as a result of ASCL1 expression, ASCL1–1 directly suppressed GCLC (Fig. S6C) resulting in a drop in cellular GSH levels. This drop in cellular GSH may be sufficient to trigger compensatory activation of the TRX pathway for cellular protection which we found to be a specific feature of NE SCLC due to ASCL1-mediated suppression of GCLC. While it has been demonstrated that breast cancer initiation is GSH-dependent while established tumors switch towards using the TRX pathway in models *in vivo (50)*, we find that non-NE SCLC maintains addiction to GSH due to the requirement to maintain GPX4 activity for protection from aberrant ferroptosis. In support of an important role for GSH during tumor initiation also in SCLC, non-NE tumors - which we find to have higher basal GSH levels-presented with a much higher tumor take-rate upon transplantation than non-NE SCLC tumors (Fig. S7A, B; non-NE 20/28 injections and for NE SCLC cells 8/28). At the same time, our data establish that NE SCLC is ferroptosis resistant but selectively sensitive to TRX pathway inhibition a novel mechanistic feature of “neuroendocrineness” in SCLC. Therefore, it will be interesting to determine whether this principle extends to other NE cancers. While several other subtype-specific therapeutic approaches have been identified for SCLC *(47, 49, 65–68)*, recent evidence suggests that SCLC subtype intratumoral heterogeneity may evolve as a consequence of subtype plasticity *(46)* highlighting the need to devise therapies targeting SCLC plasticity rather than isolated molecular subtypes. Our data from isogenic non-NE/NE cells suggest that SCLC may undergo a spontaneous switch from a GSH-dependent non-NE state towards usage of the TRX anti-oxidant pathway upon ASCL1-mediated suppression of GSH synthesis. This SCLC-specific feature of NE-non-NE plasticity may explain why, so far, the in vivo combination of BSO with Auranofin and Carboplatin has only achieved very limited efficacy in lung adenocarcinoma which is not characterized by NE-non-NE plasticity *(52)*. Supporting this notion, combined low expression of GPX4 and TXNRD1 was prognostic for drastically improved overall survival of SCLC but not LUAD patients.

Despite recent advances in the treatment of patients with SCLC, survival rates remain very poor. Our work indicates that non-NE-A SCLC is exquisitely sensitive to therapeutic induction of ferroptosis whilst NE SCLC demonstrates selective addiction to the TRX redox pathway. We anticipate that providing a novel mechanistic separation for SCLC subtype biology may on the one hand benefit patient stratification and on the other hand facilitate informed treatment approaches which take selective cell death pathway availability in SCLC subtype heterogeneity into consideration. Due to the inherent quality of lipid ROS and ROS-dependent types of cells death being molecularly independent of caspase-dependent types of cell death, we propose that this therapeutic route should remain amenable for both chemotherapy-naïve and -relapsed SCLC.

## MATERIALS AND METHODS

### Cell lines and culture conditions

Human SCLC cell lines (H82, COR-L88, H524, H889, L303, H526, H2171, H1836, H1092) were grown in suspension in RPMI (GIBCO) and were obtained from ATCC (https://www.lgcstandards-atcc.org/?geo_country=de). Murine SCLC cell lines (RP252.7, RP214, RP181.5, RP285.5, RP251, RP250) were previously derived from lung tumors of a genetically engineered mouse model for SCLC driven by loss of *Trp53* and *Rb1* by the lab of H. Christian Reinhardt. These cells were grown in RPMI. Mouse embryonic fibroblasts (MEFs) were kindly provided by Manolis Pasparakis and kept in DMEM (GIBCO), human NSCLC cell lines NCI-H460 and NCI-H727 were kindly provided by Julian Downward and kept in in DMEM (GIBCO). GPX4 KO SCLC cells were kept in RPMI with 5 μM Ferrostatin-1 (Cayman chemicals) during their generation and for part of the experiments. CDXs and PBMCs were grown in RPMI medium. All media were supplemented with 10% fetal calf serum (FCS, Sigma-Aldrich) and all cells were kept at 37°C with 5% CO_2_. All cell lines were tested for mycoplasma at regular intervals (mycoplasma barcodes, Eurofins Genomics) and human SCLC have been validated by a cell line validation service provided by Eurofins Genomics.

### Reagents

Isoleucine zipper-(iz) TRAIL was kindly provided by Henning Walczak and TNF was kindly provided by Manolis Pasparakis. The small molecules were obtained from the respective company in brackets: zVad (Enzo), Birinapant (Bertin Pharma), erastin (Biomol), RSL3 (Selleckchem), ML210 (Sigma), Ferrostatin-1 (Sigma), DFO (TargetMol), Necrostatin-1 (Abcam), NAC (BIOTREND Chemicals AG), Auranofin (Cayman Chemicals), BSO (Sigma), PX-12 (Hölzel) and D9 (Sigma). Cisplatin and etoposide were obtained from Merck. BODIPY C11 and MCB were purchased from Invitrogen and Sigma, respectively. Lipid standards (made by Avanti Polar lipids) were purchased from Sigma.

### Antibodies

Anti-Gpx4 (Abcam, ab41787, 1:2,000), Anti-xCT (Abcam, ab37185, 1:2,000), Anti-Cas9 (Cell signaling, 14697, 1:1,000,), Anti-ß-actin (Sigma, A1978, 1:10,000), Anti-GCLC (Santa Cruz, sc-166345, 1:1,000), Anti-Ascl1 (BD Pharmingen, 556604, 1:1,000), Anti-Txnrd1 (Cell signaling, 15140S, 1:1,000), Anti-Txnrd2 (Cell signaling, 12029, 1:1,000), Anti-Acsl4 (Santa Cruz Biotechnology, sc-271800, 1:2,000), Anti-Txn1 (Cell signaling, 2429S, 1:1,000), Anti-GAPDH (Cell signaling, 97166S, 1:2,000).

### Cell viability and cell death assays

Cell viability was determined by Cell Titer Blue assay (Promega) following the manufacturer’s instructions. For this assay, cells were plated at 10,000 or 5,000 cells /96-well in 100 μl media. Alternatively, cell viability was determined by Cell Titer-Glo (CTG) assay (Promega) plating 5,000 cells /96-well in 100 μl media, treatment of cells the next for 48 h followed by Cell Titer-Glo (CTG) assay (Promega) according to the manufacturer’s instructions. Cell death was quantified as cells positive for propidium iodide (PI) uptake (1 μg/ml). For this, cells were plated at 50,000 cells in 500 μl media per 24-well. Quantification was done by flow cytometry using an LSR-FACS Fortessa (BD Bioscience) counting 5,000 cells per sample.

### Generation of CRISPR/Cas9-mediated GPX4 KO cells

RP252.7 and RP285.5 were stably transfected with an expression plasmid for Cas9 (#52962 supplier) followed by selection in Blasticidin (10 μg/μl) for 5 days. Next, cells were infected with lentivirus carrying either empty vector (lentiGuide-Puro (#52963) or vector containing one of three different GPX4-targeting gRNAs and cells were kept in 5 μM Ferrostatin-1 from this time onwards:

GPX4 gRNA 1: GACGATGCACACGAAACCCC
GPX4 gRNA 2: ACGATGCACACGAAACCCCT
GPX4 gRNA 3: CGTGTGCATCGTCACCAACG

After selection with Puromycin (7 μg/ml) for 4 days whole cell populations were validated for KO via Western blot and used in Ferrostatin-1 withdrawal experiments.

### Generation of inducible Ascl1 overexpressing cells

For stable transduction of cells, viral particles were produced in HEK293T cells. First, HEK 293T cells were plated one day prior in 10 cm cell culture dish at a confluence of 70-80% at the day of transfection. Lentiviral particles were packaged using the packaging plasmids of third generation packaging system: pCMV-VSV-G (#8454) by Bob Weinberg (Stewart et al., 2003), and pRSV-Rev (#12253) and pMDLg/pRRE (#12251) by Didier Trono (Dull et al., 1998). As transfer plasmid containing the ASCL1 cDNA, pCW-Cas9 (#50661) from Eric Lander & David Sabatini, was used (Wang et al., 2013). The Cas9 fragment was firstly replaced with ASCL1 cDNA. For transfection of a 10 cm dish, 10 μg of the transfer plasmid and 5 μg of each lentiviral packaging plasmid were prepared together with 400 μl 250 mM CaCl_2_. For the formation of calcium-phosphate-DNA co-precipitate, the 400 μl of 2x HEBS buffer was added drop by drop to the CaCl_2_-DNA mixture under constant vortex. The mixture was then added drop-wise to the cells. After 6 hours transfection, cell culture medium was replaced with RPMI with 20% FCS and 1% P/S. The following three days virus containing supernatant was harvested and filtered with 0.45 μm sterile syringe filter. Target cells for transduction were plated at a confluence of 30% and virus-containing supernatant with 6 μg/ml polybrene was added on three consecutive days.

### Clonogenic survival assay

Murine SCLC cells were plated at 2,500 cells/24-well in 500 μl medium a day in advance. The next day they were treated with DMSO, RSL3 or erastin with or without Ferrostatin-1 (5 μM) for 24 h after which wells were washed with PBS and replenished with fresh media for incubation for another 6 days. On day 7, cells were washed with PBS, fixed and stained for 30 minutes using crystal violet solution (0.05% (w/v) crystal violet, 1% Formaldehyde, 1% methanol in PBS).

### Western blotting

Cells were washed, lysed in IP-lysis buffer (30 mM Tris-HCl, 120 mM NaCl, 2 mM EDTA, 2 mM KCl, 1% Triton-X-100, pH 7,4, Protease and Phosphatase inhibitor (Roche) and frozen at −20°C. After re-thawing, lysate concentrations were adjusted to equal protein concentrations using the bicinchoninic acid (BCA) protein assay (Biorad). Equal amounts of protein were mixed to a final concentration of 1x reducing sample buffer (Invitrogen) containing 200 mM DTT. Samples were heated to 80°C for 10 minutes, separated via gel electrophoresis and transferred to PVDF membranes using the TurboBlotting system (Biorad). Membranes were blocked in PBS with 0.1% Tween 20 (PBST) with 5% (w/v) dried milk powder for at least 30 minutes. Next, membranes were incubated over night at 4°C with primary antibodies in PBST with 5% bovine serum albumin (BSA). After washing with PBST, membranes were incubated with horse radish peroxidase (HRP)-coupled secondary antibodies for at least 1h. After another washing step, membranes were developed using chemiluminescent substrate Immobilon Luminata Classico (Millipore) and X-ray films CL-XPosure™ (Thermo Scientific).

### Lipid ROS quantification by BODIPY C11 staining

For lipid ROS quantification, 50,000 cells were seeded in 500 μl media in 24-well plates. The next day, cells were treated with DMSO, 100 nM RSL3 or 100 nM RSL3/5 μM Fer-1 for 5 h. During the last 30 minutes of incubation, BODIPY C11 was added at 5 μM to each well. Cells were then washed, detached and increased green fluorescence over baseline (stained but untreated) was determined by flow cytometry (excitation 520 nm) counting 5,000 cells per sample.

### Cellular GSH quantification

To determine relative levels of GSH in responders and non-responders, the thiol-reactive dye monochlorobimane (MCB) was added to cells seeded a day in advance (50,000 cells per well in 24-well plates) at a concentration of 50 μM for 30 minutes. Cells were then washed, detached and analyzed by flow cytometry (405 nm excitation) counting 5,000 cells per sample.

To quantify concentrations of cellular GSH and GSSG, the GSH/GSSG Glo Assay (Promega) was used according to the manufacturer’s instructions using 75,000 (H82 and H2171) or 50,000 (RP) cells.

### ASCL1-staining for Flow Cytometry

RP181.5 stickers or floaters (1×10^5^ cells) were seeded in 12-well plates a day in advance and then treated with 5 mM BSO or 1 μM Auranofin for 24 h. Cells were harvested, washed with PBS and stained for live/dead cells using the viability dye eFluor780 (eBioscience), 1:1000), for 30 min, at 4°C. Cells were then washed twice with FACS buffer (PBS, 2% FCS) and cell pellets were resuspended in 200 μl Fixation/Permeabilization buffer (eBioscience) (overnight incubation at 4°C). The next day, cells were washed with 1x Permeabilization buffer (eBioscience) and incubated for 15 min in FACS buffer before adding the primary MASH-1 antibody (BD Biosciences, 556604), 1:250, for 30 min at 4°C in 1x Permeabilization buffer. After washing cells twice with 1x Permeabilization buffer, pellets were resuspended in the secondary antibody (Cy3, Jackson Laboratories) 1:500 for 30 min at 4°C in 1x Permeabilization buffer. Cells were again washed twice with 1x Permeabilization buffer and resuspended in FACS buffer. Measurements were acquired using a BD LSR Fortessa flow cytometer and data were analyzed with the FlowJo software. For sorting experiments cells were resuspended in PBS with 2% FCS and 25 mM Hepes and passed through a BD FACS sorter (Influx).

### Redox shift assays

1.5×10^6^ cells (H82, RP181.5 floaters) and 1×10^6^ cells (H2171, RP181.5 stickers) were seeded in 6-well plates and treated for 0.5 or 6 h with 1 μM Auranofin. Cells were then harvested, washed in PBS and lysed in 1 ml 8% (w/v) trichloroacetic acid (TCA) on ice and frozen at −20°C. Samples were thawed and centrifuged for 15 min at 20,000 g at 4°C. TCA supernatant was removed, pellets were centrifuged for 5 min at 20,000 g at 4°C and residual TCA was removed. 20 μl of alkylation buffer (6 M urea, 0.2 M Tris-HCl, pH 7.5, 0.2 M EDTA, 2% SDS, bromophenol blue) was added and samples were sonicated for 10 cycles at an amplitude of 60% to dissolve pellets (UP50H, Hielscher). For minimum and maximum samples, tris(2-carboxyethyl)phosphine (TCEP) was added (10 μM final concentration) followed by incubation for 10 min at 50°C. For the minimum shift sample, N-ethyl maleimide (NEM) was added (15 mM final concentration) and to all other samples methyl-polyethylenglycol-maleimide (mmPEG24, 15 mM final concentration) was added. Samples were incubated for 1 h in the dark at room temperature, then 20 μl of 2x Laemmli buffer was added. Samples were loaded on tris-tricine gels and run overnight at constant 130 V at 4°C.

### Quantification of oxidized glycerophospholipids via lipidomics

Levels of oxidized phosphatidylcholine (PC) and phosphatidylethanolamine (PE) species were determined by Liquid Chromatography coupled to Electrospray Ionization Tandem Mass Spectrometry (LC-ESI-MS/MS) using a procedure described in Doll et al. 2019 *(22)* with several modifications: 2.5×10^6^ cells were resuspended in 250 μl of an ice-cold 2:5 (v/v) mixture of 100 μM diethylenetriaminepentaacetic acid (DTPA) in PBS, pH 7.4, and 40 μM butylated hydroxytoluene (BHT) in methanol. To 100 μl of the cell suspension another 3.4 ml of the above-mentioned PBS/methanol mixture, 1.25 ml of ice-cold chloroform and internal standards (10 pmol 1,2-dimyristoyl-*sn*-glycero-3-phosphocholine (DMPC) and 10 pmol 1,2-dimyristoyl-*sn*-glycero-3-phosphoethanolamine (DMPE) were added. The samples were vortexed for 1 min and incubated at −20°C for 15 min. After adding 1.25 ml of chloroform and 1.25 ml of water, the mixture was vortexed vigorously for 30 sec and then centrifuged (4,000 × g, 5 min, 4°C) to separate layers. The lower (organic) phase was transferred to a new tube and dried under a stream of nitrogen. The residues were resolved in 150 μl of methanol and transferred to autoinjector vials. LC-MS/MS analysis was performed by injecting 20 μl of sample onto a Core-Shell Kinetex C18 column (150 mm × 2.1 mm ID, 2.6 μm particle size, 100 Å pore size, Phenomenex) at 30°C and with detection using a QTRAP 6500 triple quadrupole/linear ion trap mass spectrometer (SCIEX). The LC (Nexera X2 UHPLC System, Shimadzu) was operated at a flow rate of 200 μl/min. The mobile phase system, gradient program and source- and compound-dependent parameters of the mass spectrometer were set as previously described *(22)(22)(20)*(Doll et al., 2019). Oxidized PC and PE species and the internal standards were monitored in the negative ion mode using their specific Multiple Reaction Monitoring (MRM) transitions *(22)*. The LC chromatogram peaks were integrated using the MultiQuant 3.0.2 software (SCIEX). Oxidized PC and PE species were quantified by normalizing their peak areas to those of the internal standards. The normalized peak areas were related to the mean values of the *floaters* + DMSO cell samples.

### IncuCyte cell imaging

Cells were plated in 96-well plates (1× 10^4^ *stickers*, 2 × 10^4^ *floaters*) and stimulated with or without Ferrostatin-1 [5μM] 2 h prior to adding either DMSO, BSO [10mM] or Auranofin [1μM]. Dead cells were stained by adding 100 nM DRAQ7 (Thermofisher) to all wells. Cells were imaged for 24 h every 5 h and 3 images per well were captured.

### ASCL1-staining for immunofluorescence microscopy

RP181.5 stickers (5000 cells) were seeded in 24-well plates. The following day, they were treated with 500 nM Auranofin and 500 μM BSO for 96 h. For staining cells were washed with PBS followed by incubation in 200 μl Fixation/Permeabilization buffer (eBioscience) overnight at 4°C. The next day, cells were washed with 1x Permeabilization buffer (eBioscience) and incubated for 15 min in FACS buffer (PBS, 2% FCS) before adding the primary MASH-1 antibody (BD Biosciences, 556604), 1:250, for 30 min at 4°C in 1x Permeabilization buffer. After washing cells twice with 1x Permeabilization buffer, cells were incubated in the secondary antibody (Cy3, Jackson Laboratories) 1:500 for 30 min at 4°C in 1x Permeabilization buffer. Cells were again washed twice with 1x Permeabilization buffer and subsequently incubated in PBS for microscopy. Microscopy was performed using a Leica DMI 6000B microscope (20x objective) and images were analyzed with the ImageJ software.

### Protein isolation from fresh frozen tissue

Fresh frozen tumor tissue was weighed and mixed in a peqlab vial with an adequate amount of ceramic beads and 500 μl lysis buffer (30 mM Tris-HCl, 120 mM NaCl, 2 mM EDTA, 2 mM KCl, 1% Triton-X-100, pH 7,4, Protease Inhibitor (Roche). For lysis, samples were homogenized for 2 x 30 sec using the Precellys 24-dual homogenisator (Peqlab) and incubated on ice for 30 min.

### Tumor xenograft studies

1.5×10^6^ cells (either H82 or H2171) in 200 μl PBS were injected into both flanks of 8-week old male NMRI-Foxn1 nu/nu mice (Janvier). Mice were enrolled either into vehicle or combination treatment groups once tumors reached a minimum size of 2 × 2 mm. For two consecutive weeks mice were injected either with vehicle (PBS with 2% DMSO, 8.5% ethanol and 5% polyethylene glycol 400) and received normal drinking water or were injected with Auranofin (2.5 mg/kg) three times a week and received BSO (5 mM) in the drinking water *ad libitum*. Tumor size was tracked by caliper measurements and volume was calculated as (length*width*width)/2. Mice were sacrificed at the end of the treatment. Due to variance in tumor size at the start of the experiment, fold change of tumor volume upon treatment is plotted. All animal experiments were approved by the local authorities (LANUV, North-Rhine-Westphalia, Germany) and performed under license number 81-02.04.2017.A477. All people involved in animal experiments received prior training and have passed the additionally required personal licensing course (FELASA-B). All animal experiments were conducted in compliance with international and institutional ethical guidelines on animal welfare and measures to minimize animal suffering.

### Immunohistochemistry

For immunohistochemistry tissues were fixed in 4% Formaldehyde solution (Merck) and subsequently embedded in paraffin. After sectioning tissue (4 μm) it was deparaffinized and rehydrated according to standard procedure. For antigen retrieval samples were heated in a citrate-based buffer (Vector Laboratories, H-3300) to 100°C for 30 min. Subsequently, samples were blocked for endogenous peroxidases (BLOXALL Endogenous Peroxidase and Alkaline Phosphatase Blocking Solution, Vector Laboratories, SP-6000, 15 min), unspecific bindings (5% BSA, 5% NGS in PBST, 1h) and Avidin/Biotin (Avidin/Biotin Blocking Kit, Vector Laboratories, SP-2001, 15 min). Samples were incubated overnight at 4°C with anti-MDA antibody (Abcam, ab6463) 1:250 in blocking buffer (PBS, 1% BSA, 0.003% NaN3, 0.05% Tween20). The following day samples were incubated with secondary antibody (Perkin Elmer, NEF813) 1:1,000 in blocking buffer for 1 h. All blocking steps were followed by three PBS washing steps. For staining samples were incubated with PBS + 1/60 Biotin + 1/60 Avidin (VECTASTAIN® Elite® ABC HRP Kit, Vector Laboratories, PK-6100) for 30 min followed by DAB staining (Abcam, ab6423) according to the manufacturer’s instructions.

### RP-mouse model, MRI scans, treatment

The well-established genetically engineered mouse model recapitulating SCLC harbors a Rb1flox/flox allele in which exons 18 and 19 are flanked by loxP sites as well as a Trp53flox/flox allele in which exons 2-10 are flanked by loxP sites (RP model). RP mice used in this experiments were kept on a mixed C57Bl6/J;Sv129 background. For induction of lung tumor formation, 8-12 weeks old mice were anesthetized with Ketavet (100 mg/kg) and Rompun (20 mg/kg) by intraperitoneal injection followed by intratracheal inhalation of replication-deficient adenovirus expressing Cre (Ad5-CMV-Cre, 2.5×107 PFU, University of Iowa) to their lungs *(41)*. Five months after tumor induction, tumor formation was monitored bi-weekly by magnetic resonance imaging (MRI) (A 3.0 T Philips Achieva clinical MRI (Philips Best, the Netherlands) in combination with a dedicated mouse solenoid coil (Philips Hamburg, Germany), were used for imaging. T2-weighted MR images were acquired in the axial plane using turbo-spin echo (TSE) sequence [repetition time (TR)=3819 ms, echo time (TE)=60 ms, field of view (FOV)=40×40×20 mm^3^, reconstructed voxel size=0.13×0.13×1.0 mm^3^, number of average=1) under isoflurane (2.5%) anesthesia. MR images (DICOM files) were analyzed by determining and calculating region of interests (ROIs) using Horos software. Once tumors reached a mean volume of 15-30 mm^3^, mice were randomized into two groups and treated with either vehicle or the combination of BSO (5 mM, via drinking water) and Auranofin (2.5 mg/kg, 3x per week, i.p.) for 2 weeks. All animal experiments were approved by the local authorities (LANUV, North-Rhine-Westphalia, Germany) and performed under license number 81-02.04.2017.A477. All people involved in animal experiments received prior training and have passed the additionally required personal licensing course (FELASA B). All animal experiments were conducted in compliance with international and institutional ethical guidelines on animal welfare and measures to minimize animal suffering.

### Generation of human SCLC CDXs

Circulating tumor cells (CTCs) were isolated from blood of two patients diagnosed with SCLC following previously described protocols *(54)*. After the development of a tumor in immunocompromised NSG mice, cells were dissociated, expanded and maintained in cell culture in HITES media. At least 90% of cells were confirmed to be tumor cells with neuroendocrine marker expression (chromogranin A, CD56 and synaptophysin) as determined by RNA analysis and immunohistochemistry (IHC). Use of patient material was approved by the institutional review board of the University of Cologne following written informed consent. We have complied with all relevant ethical regulations pertaining to the use of human patient material.

### PBMC isolation

PBMCs were isolated from buffy coats from two healthy donors under an existing ethics approval at the University Hospital Cologne (01-090) following written informed consent. We have complied with all relevant ethical regulations pertaining to the use of human patient material.

### Human SCLC RNA-seq and survival data availability

All SCLC and normal lung patient datasets and human SCLC cell line data used in this study have been previously published and, as such, are appropriately referenced, have previously obtained the appropriate ethics approvals and are available online under https://doi.org/10.1038/nature14664 for George et al. 2015 (SCLC data), https://doi.org/10.1038/ng.2405 for Rudin et al. 2012 (normal lung data) and under https://doi.org/10.1016/j.ccell.2016.12.005 for Mollaoglu et al. 2017 (human SCLC cell line RNA-seq data). Raw sequencing data were analyzed with TRUP *(69)* and gene expression was quantified as FPKM.

### Analysis software and bioinformatic analysis

Heatmaps vizualizing cell death pathway component expression were generated using RStudio version 1.1.456 and gplots and RColorBrewer packages. A ranked list of fold differential expression was generated for human cell line RNA-seq data using Excel and analyzed by GSEA Desktop v3.0 (https://doi.org/10.1073/pnas.0506580102 and https://doi.org/10.1038/ng1180). FACS data were analyzed and quantified using FlowJo 10.4.2. Cell Titer Blue viability assays were analyzed using Excel. MRI scans were quantified using Horos v3.3.5. Lipidomics measurements were analyzed by MultiQuant 3.0.2 software (SCIEX). Figures were assembled and data plotted and analyzed using GraphPad Prism 7 for Mac OS X.

### Data presentation and statistical analysis

Data are presented as mean +/− SEM of at least three independent experiments unless stated otherwise. Thereby, the mean is calculated and plotted from at least three means from three independent experiments (each one performed in duplicates or triplicate experimental replicates). Two-tailed p values with a cut-off of *=0.05 are calculated for the comparison between two conditions/groups. For comparison of multiple conditions/groups p values (cut off *=0.05) are determined by ANOVA and Bonferroni post-test. All statistical analysis was performed using GraphPad Prism 7 for Mac OS X.

## Supporting information

Bebber et al Supl. Data

## SUPPLEMENTARY MATERIALS

Fig. S1. SCLC is more prone to respond to erastin than NSCLC.

Fig. S2. SCLC is sensitive to induction of ferroptosis.

Fig. S3. SCLC divides into ferroptosis responders and non-responders.

Fig. S4. Chemotherapy triggers caspase-dependent cell death in SCLC.

Fig. S5. NE-A SCLC is selectively resistant to ferroptosis and presents with low ASCL4 expression.

Fig. S6. NE-A SCLC suppresses cellular GSH levels and creates dependency on the thioredoxin pathway.

Fig. S7. BSO/Auranofin-induced cell death is partially ferroptotic and partially dependent on ROS.

Table S1. Supplementary Table 1.

## Acknowledgments

We thank Karin Schlegelmilch, Snehlata Kumari and all members of the Department of Translational Genomics for constructive feedback, Henning Walczak and Manolis Pasparakis for providing recombinant TRAIL and TNF, respectively, and Johannes Kühle for providing buffy coats.

## Author contributions

C.M.B. performed many in vitro and the majority of in vivo experiments, designed experiments and supervised Z.C. and L.S., E.S.T. performed the majority of the in vitro experiments and bioinformatic analysis, Z.C. and J.S. performed knockdown experiments and experimental repeats, A.S. assisted with RP mouse in vivo experiments, A.A. performed redox pathway plasticity FACS experiments, M.N.H. performed redox shift assays, L.S. assisted with tissue stainings, M.A.D. performed human SCLC cross titrations, C.P.A. cultured primary CDXs, A.K. performed human cell line RNA-seq data analysis, F.P. assisted with human SCLC cross titrations, F.B. performed TCGA LUAD survival analysis, M.L.S. designed human SCLC cross titration experiments, J.R. designed redox shift experiments, J.G. provided CDXs and gene expression data, S.B. performed lipidomics measurements, R.K.T and H.C.R. designed experiments and S.v.K. conceived the project, designed experiments, supervised work, acquired funding and wrote the manuscript. All authors read and edited the manuscript.

## Data and materials availability

The authors acknowledge that part of the RNA-seq data were generated by Genentech/gRED, previously published by Rudin et al. *(70)*.

