## Supplementary material for "Ferroptosis response segregates small cell lung cancer (SCLC) neuroendocrine subtypes": Bebber et al Supl. Data

**Supplementary Materials: Bebber et al.**

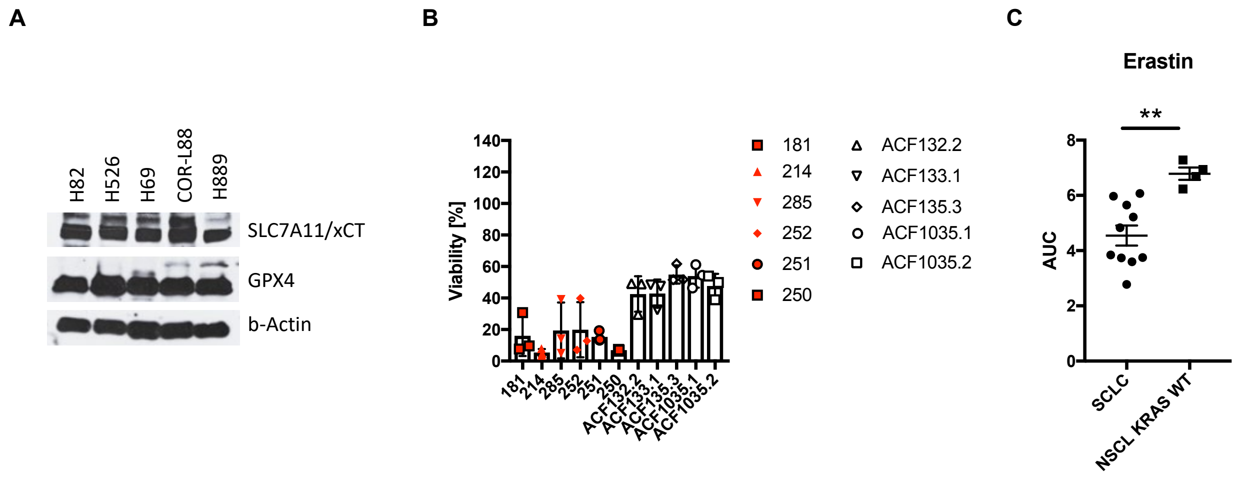


**Figure S1. SCLC is more prone to respond to erastin than NSCLC.** (**A**) western blot of ferroptosis pathway component expression in representative human SCLC cell lines. (**B**) the indicated murine SCLC (red; n=6) and NSCLC (white; n=5) cell lines were treated with erastin [10 µM] for 24 h, cell viability was determined by Cell Titer Blue. (**C**) erastin-treated human SCLC (n=10) and KRAS wild type NSCLC cells (n=4) from Yang et al. *(18)* were plotted for area under the curve (AUC). Data are means +/- SEM of three independent experiments for each cell line wherever not indicated otherwise.

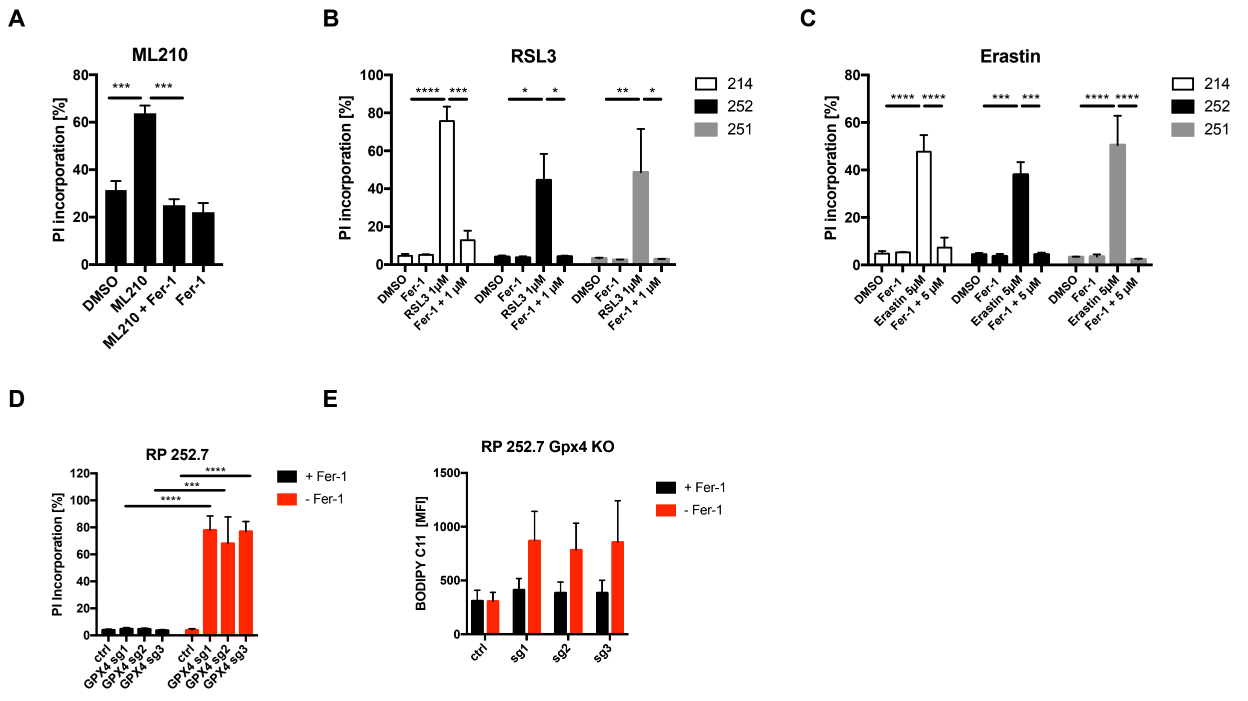


**Figure S2. SCLC is sensitive to induction of ferroptosis.** (**A**) Human H82 cells were with ML210 [1 µM] +/- Fer-1 for 24 h. Cell death was determined by propidium iodide (PI) uptake and flow cytometry. (**B**, **C**) the indicated murine SCLC cells were treated as indicated erastin [10 µM] +/- Ferrostatin-1 (Fer-1, 5 µM) +/- Deferoxamine (DFO) [100 µM], RSL3 [1 µM] +/- Fer-1 +/- DFO for 24 h and quantified as in (A). Data are means +/- SEM of three independent experiments. **(D)** RP252.7 cells with either control or GPX4-targeting gRNA were grown in the presence of 5 µM Fer-1. Cell death was determined after 24 h of Fer-1 withdrawal by propidium iodide (PI) uptake and flow cytometry. (**E**) cells, as in a, were cultured in the presence or withdrawal of Fer-1 [5 µM] for 5 h and stained for lipid ROS accumulation using BODIPY C11. Cells were analyzed by flow cytometry and mean fluorescent intensity (MFI) was quantified.

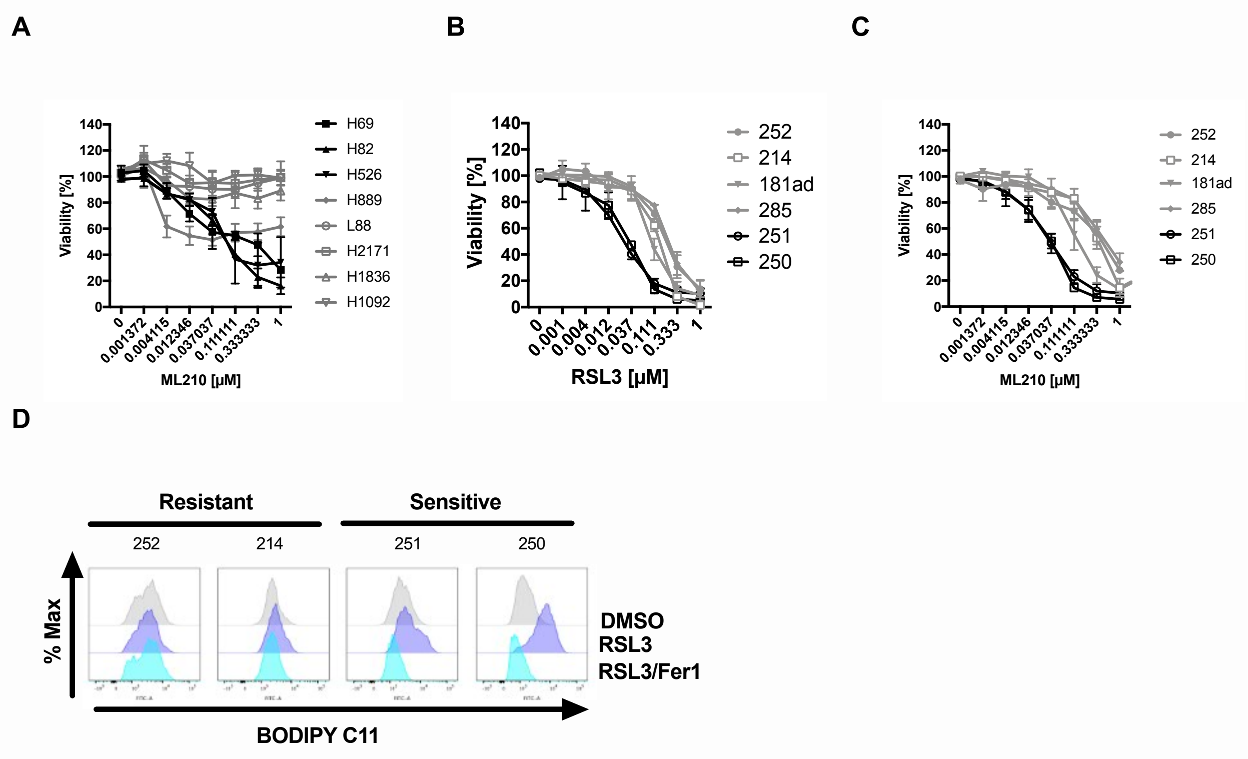


**Figure S3. SCLC divides into ferroptosis responders and non-responders.** (**A**) the indicated human SCLC cell lines (n=8) were treated as indicated for 24 h, cell viability was determined by Cell Titer Blue. (**B**) murine SCLC cell lines (n=6) were treated as indicated for 24 h, cell viability was determined by Cell Titer Blue. Data are means +/- SEM of three independent experiments in each individual cell line or representative images were applicable. (**C**) murine SCLC cell lines (n=6) were treated as indicated for 24 h, cell viability was determined by Cell Titer Blue. (**D**) indicated murine SCLC cells were treated with DMSO, RSL3 [300 nM] +/- Fer-1 [5 μM] for 5 h and stained for lipid ROS accumulation using BODIPY C11. Cells were analyzed by flow cytometry. Data are means +/- SEM of three independent experiments or representative FACS histograms where applicable.

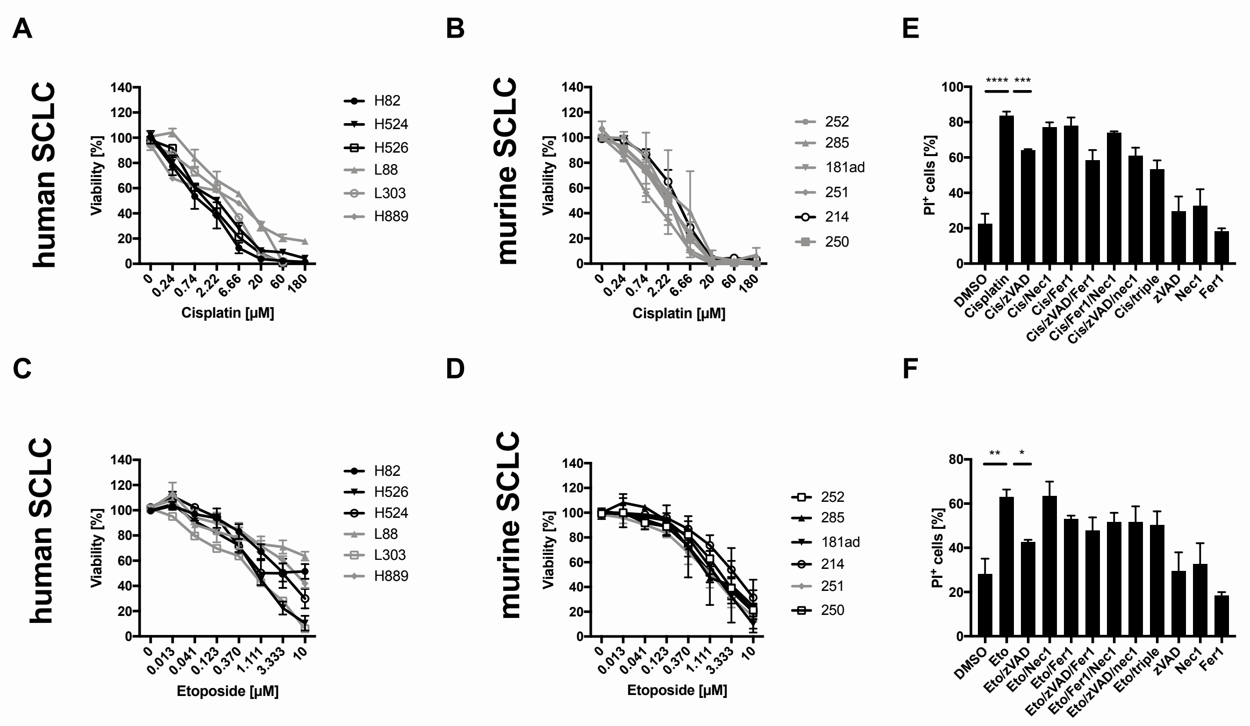


**Figure S4. Chemotherapy triggers caspase-dependent cell death in SCLC.** (**A**, **C**) indicated human SCLC lines (n=7) were treated with rising concentrations of cisplatin or etoposide for 72 h, cell viability was determined by Cell Titer Blue. (**B**, **D**) indicated murine SCLC lines (n=6) were treated as indicated for 72 h, cell viability was determined by Cell Titer Blue. (**E**, **F**) H82 cells were treated with DMSO, cisplatin [180 µM] or etoposide [10 µM] and the indicated combinations using zVAD [20 µM], necrostatin-1 (Nec1) [10 µM] or Ferrostatin-1 (Fer-1) [5 µM] for 72 h. Cell death was determined by propidium iodide (PI) uptake and flow cytometry. triple, zVAD/Nec1/Fer-1; Data are means +/- SEM of three independent experiments.

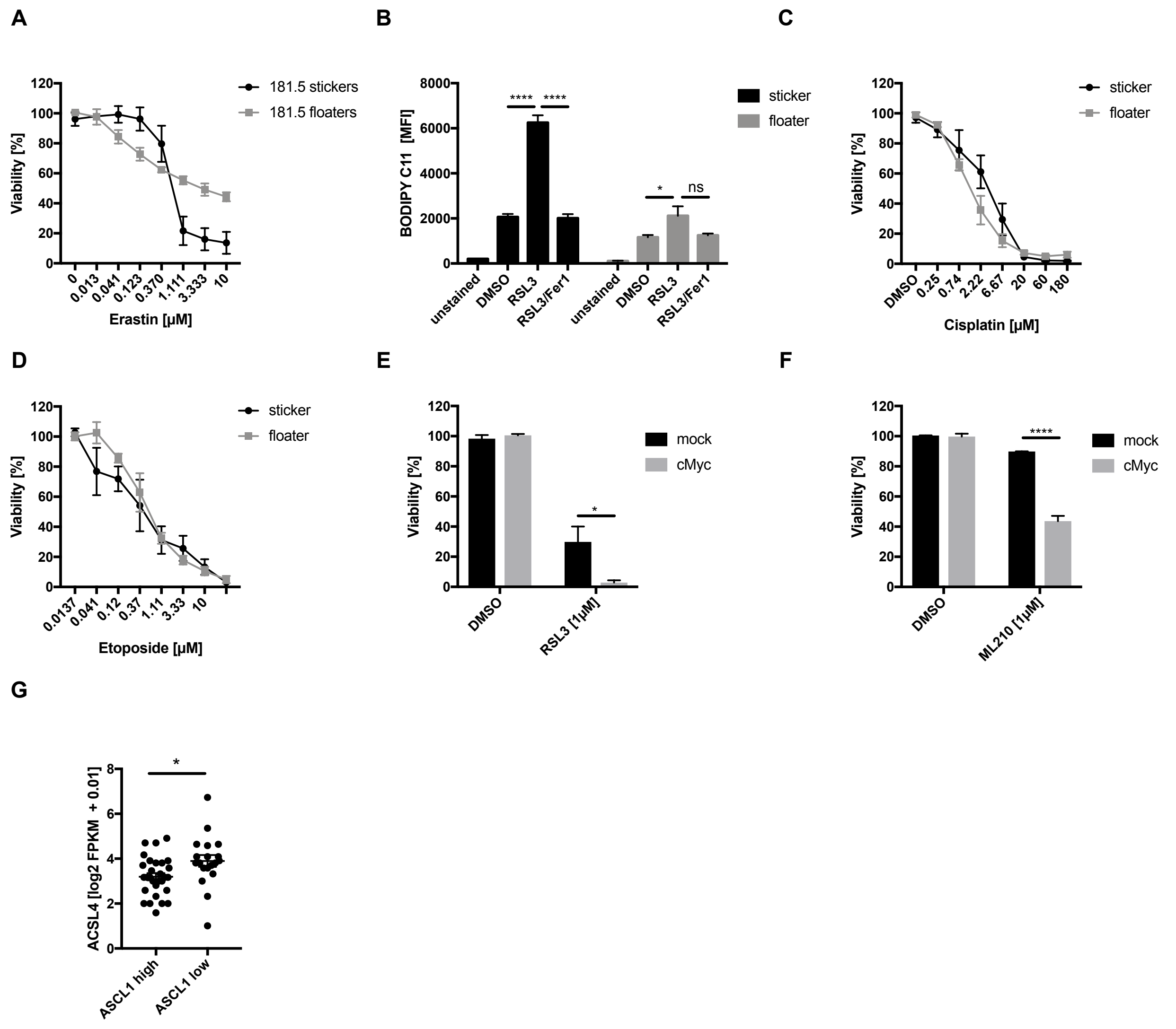


**Figure S5. NE-A SCLC is selectively resistant to ferroptosis and presents with low ASCL4 expression.** (**A**) *stickers* and *floaters* were cultured separately and treated with erastin at the indicated concentrations for 24 h, cell viability was determined by Cell Titer Blue. (**B**) cells as in (**A**) were treated with DMSO, RSL3 [1 µM] +/- Fer-1 [5 µM] for 5 h and stained for lipid ROS accumulation using BODIPY C11 and analyzed by flow cytometry; mean fluorescent intensity (MFI) was quantified. (**C, D**) cells as in a were treated with rising concentrations of cisplatin or etoposide for 72 h, cell viability was determined by Cell Titer Blue. (**E**, **F**) murine RP cells expressing cMyc from the endogenous locus via CRISPRa *(49)* were treated as indicated for 24 h, cell viability was determined by Cell Titer Blue. (**G**) the NIH SCLC cell line panel (n=48) was segregated by ASCL1 expression (high= log2 (FPKM +0,01)> 3) (low= log2 FPKM +0,01< 3)

and ACSL4 mRNA expression is plotted. Cell line panel expression data are available at Expression Atlas https://www.ebi.ac.uk/gxa/home. Data are means +/- SEM of three independent experiments or representative images where applicable.

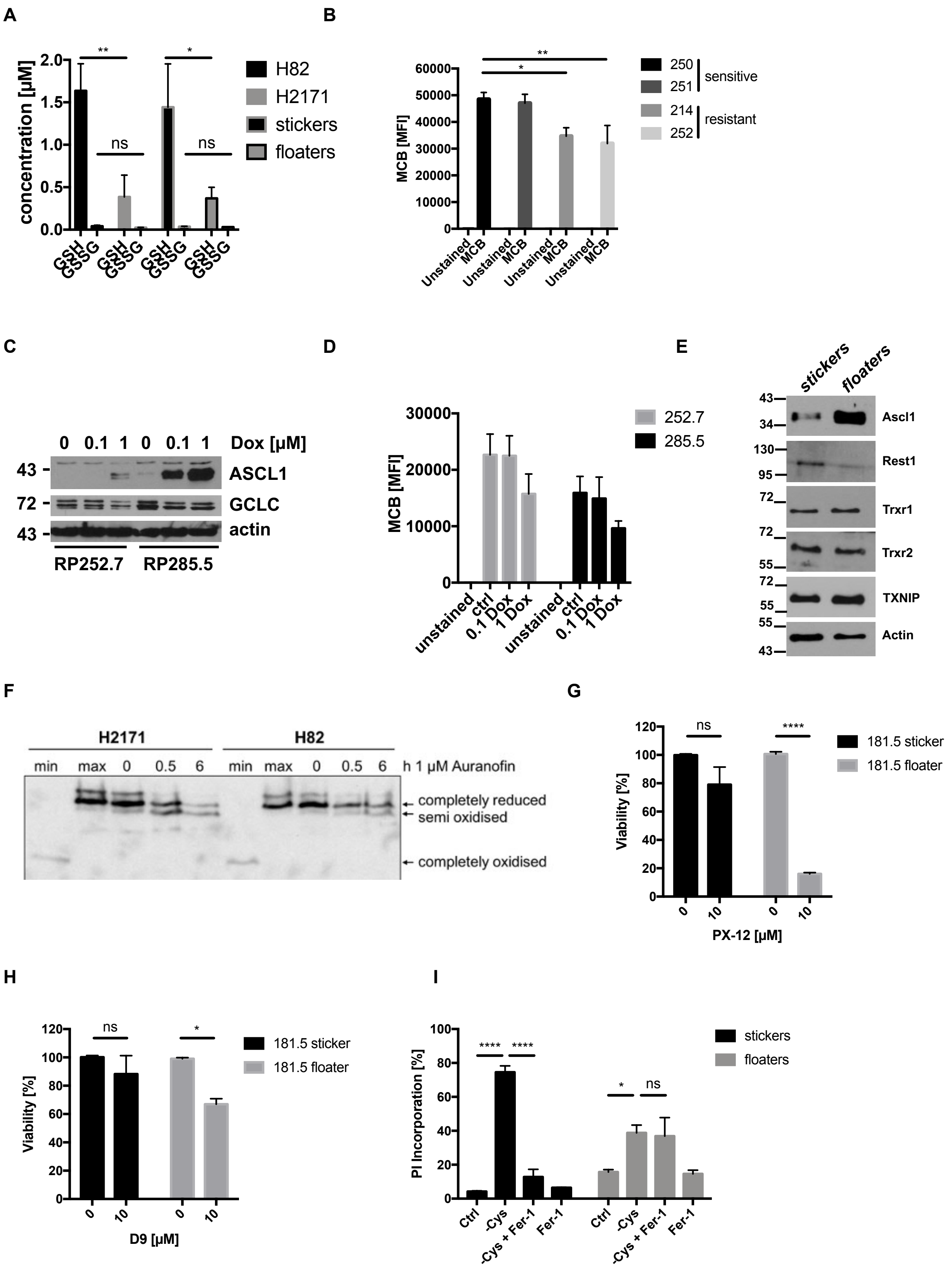


**Figure S6. NE-A SCLC suppresses cellular GSH levels and creates dependency on the thioredoxin pathway.** (**A**) Cellular GSH concentrations were quantified by GSH/GSSG Glo

Assay (Promega) in the indicated cell lines. (**B**) cellular GSH levels were determined by monochlorobimane (MCB) staining and flow cytometry (MFI- mean fluorescent intensity) in the indicated murine SCLC cell lines. (**C**) the indicated ASCL1-inducible cell lines were treated as indicated for 24 h. Representative western blots are shown. (**D**) cellular GSH levels were determined by monochlorobimane (MCB) staining and flow cytometry (MFI- mean fluorescent intensity) in the indicated murine SCLC cell lines upon 24h of doxycline induction. (**E**) *stickers* and *floaters* were cultured separately and subjected to Western Blot analysis. (**F**) $1.5x10^6$ (H82) and $1x10^6$ (H2171) cells were treated with Auranofin [1 μM] for the indicated times, cells were lysed with 8% (w/v) TCA. TRX shift was analyzed by Western Blot. (**G**, **H**) *stickers* and *floaters* were cultured separately and treated as indicated for 48 h. Cell viability was determined by Cell Titer Blue. (**I**) *stickers* and *floaters* were cultured in normal (Ctrl) or cystine-free medium (-Cys) +/- Fer-1 [5 μM] for 24 h, cell death was quantified by propidium iodide (PI) uptake and flow cytometry. Data are means +/- SEM of three independent experiments or representative Western blots where applicable.

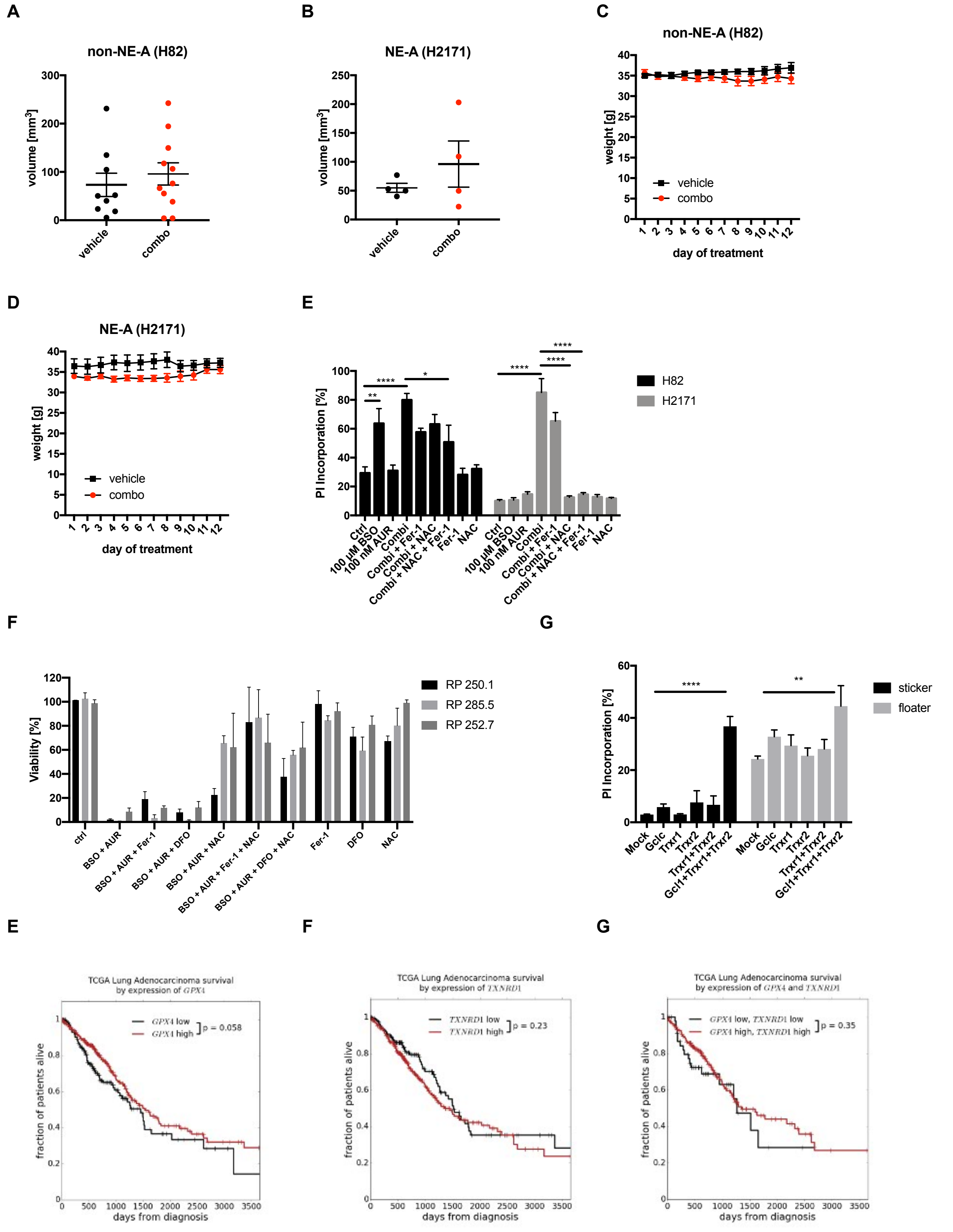


**Figure S7. BSO/Auranofin-induced cell death is partially ferroptotic and partially dependent on ROS** (**A, B**) 8-weeks old male nude mice were injected with $1.5x10^6$ H82 (n=20 tumors) or H2171 (n=8 tumors) cells into flanks. Once measurable, tumors were randomized for the indicated treatment groups. (**C, D**) mice were treated either with vehicle or combined BSO [5 mM] in the drinking water and Auranofin 3x per week intraperitoneal (i.p.) [2.5 mg/kg] for two consecutive weeks. Mouse weights were tracked during this time. (**E**) the indicated human SCLC cell lines were treated with the indicated combinations of BSO [10 mM], Auranofin (AUR) [1 µM], N-Acetyl Cysteine (NAC) [3 mM], Ferrostatin-1 (Fer-1) [5 µM] for 24 h. Cell death was quantified by propidium iodide (PI) uptake and flow cytometry. (**F**) the indicated murine SCLC cell lines were either left untreated (ctrl), or treated as indicated: BSO [10 mM], Auranofin (AUR) [1 µM], Deferoxamine (DFO) [100 µM], N-Acetyl Cysteine (NAC) [3 mM], Ferrostatin-1 (Fer-1) [5 µM] for 24 h. Cell viability was determined by Cell Titer Blue. (**G**) *stickers* and *floaters* were subjected to siRNA-mediated knockdown targeting the indicated proteins for 72 h. Spontaneous cell death was quantified by propidium iodide (PI) uptake and flow cytometry. Data are means +/- SEM or representative pictures out of three independent experiments. (**H**) Kaplan-Meier survival curves for LUAD patients from TCGA (n=503) containing low (low 1/3 n=167, median survival 48.5 months) or high (high 2/3 n=336, median survival 51 months) expression of GPX4 mRNA. (**I**) as in (**H**), expression of TXNRD1 mRNA was correlated using the same cut-off (low=1/3, median survival 50.5 months; high 2/3, median survival 45.2 months). (**J**) Kaplan-Meier survival curves for LUAD patients from TCGA with combined low or high GPX4 and TXNRD1 mRNA expression (low/low n=49, median survival 42.2 months; high/high n=218, median survival 45.2 months).

**Additional Files:**

Supplementary Table 1: Lipidomics Data
